# Notch2 signaling guides B cells away from germinal centers towards marginal zone B cell and plasma cell differentiation

**DOI:** 10.1101/2022.06.13.495961

**Authors:** T. Babushku, M. Lechner, A. J. Yates, S. Rane, U. Zimber-Strobl, L. J. Strobl

## Abstract

Notch2 signaling has a profound role in driving the development of Marginal Zone B (MZB) cells. We recently demonstrated that Follicular B (FoB) cells act as precursors for MZB cells in mice, but the mechanistic aspects of this differentiation pathway are still elusive. By studying Notch signaling in CBF:H2B-Venus Notch-reporter mice, we show that most B cells receive a Notch signal, which is highest in MZB cells. However, surprisingly, around one-third of MZB cells seem to lose their Notch signal with time. Conditional deletion or constitutive activation of Notch2 in mice upon T-cell-dependent (TD) immunization unraveled an interplay between antigen-induced activation and Notch2 signaling, in which FoB cells that turn off the Notch pathway enter germinal centers, whereas FoB cells with high Notch signals undergo MZB cell or plasmablast differentiation. Input of experimental data into a mathematical modeling framework reveals that MZB cells regularly emerge from antigen-activated FoB cells in a Notch2-dependent manner upon TD immunization.

## Introduction

Mature splenic B2 cells are comprised of two distinct cell populations: Follicular B (FoB) and Marginal Zone B (MZB) cells. FoB cells recirculate between lymphoid organs, whereas MZB cells are mostly sessile in the marginal zone (MZ) and can occasionally migrate between the B cell follicle and the MZ (1). MZB cells respond to T-cell-independent (TI) antigens by differentiation into low affinity IgM-producing plasma cells (PCs) (2, 3). In contrast, FoB cells respond to T-cell-dependent (TD) antigens, giving rise to germinal centers (GC), where they undergo class switching and somatic hypermutation, resulting in the generation of high-affinity Ig-switched PCs and memory B cells (4, 5).

B cell development begins in the bone marrow (BM) and is completed in the spleen. Immature B cells leave the BM upon successful rearrangement of the B-cell-receptors (BCR), and pass through short-lived developmental stages (T1-T3) before differentiating into either FoB or MZB cells in the spleen (6–8). MZB cells may also develop from FoB cells, in particular FoB-II cells, a recirculating long-lived population with an IgM^high^IgD^high^CD21^int^CD23^+^ phenotype (9–13). We recently strengthened this hypothesis by showing that the adoptive transfer of highly purified FoB cells into immunocompetent recipients led to the generation of donor-derived MZB cells via an intermediate MZ precursor stage (14). Our data provided strong evidence that FoB cells can act as precursors for MZB cells and we postulated that this could be physiologically mediated via activation of Notch2.

Notch2 signaling has been long known to be a crucial player in MZB cell development. Constitutively active Notch2 signaling in T1 or FoB cells strongly induces MZB cell differentiation (14, 15). In contrast, homozygous or heterozygous inactivation of Notch2 in B lymphocytes or of its ligand Dll-1 in non-hematopoietic cells leads to the total loss or reduction of MZB cells, respectively (16, 17), suggesting a dependence of MZB cell development on ligand/receptor dosage. This phenomenon may imply that the bottleneck in MZB cell development is the extent of Notch2 signaling, which is modulated by Notch2 surface expression and its modification (16, 18, 19) as well as the availability of Dll-1-expressing fibroblasts (20). Additionally, BCR-signaling strength has been assumed to influence MZB cell fate decisions (6, 21). We have shown that Notch2 surface expression is strongly upregulated by BCR-stimulation *in vitro* (14) and others found that BCR-signaling induces recruitment of Adam10 to the plasma membrane, enhancing the cleavage and subsequent activation of Notch2 (22). These findings suggest that BCR signals act upstream of Notch2 signaling and in combination with CD40 signals, increase the likelihood of B cells to acquire a strong Notch2 signal that drives their commitment to the MZB cell fate.

By studying Venus expression in different B cell populations of Notch-reporter mice, we now demonstrate that almost all B cells receive a Notch signal under steady-state conditions, and this signal is strongest in MZB cells. Moreover, we provide evidence that induction of Notch2 signaling upon TD immunization navigates cells away from the GC reaction and induces PC and MZB cell differentiation instead, thus making the Notch2 signaling pathway an important binary switch between opposed B cell fates in the course of TD immune responses.

## Results

### All B cells receive a Notch signal under steady-state conditions

To identify the B cell populations that receive a Notch signal, we analyzed B lymphocytes in the BM and spleen of CBF:H2B-Venus mice, which express Venus in all cells that undergo active or have recently experienced Notch signaling (23). We found that almost all developing B cell populations in the BM were Venus^+^, with recirculating B cells (CD43^-^B220^high^) having the highest Venus expression (Fig. 1A).

**Fig. 1.**
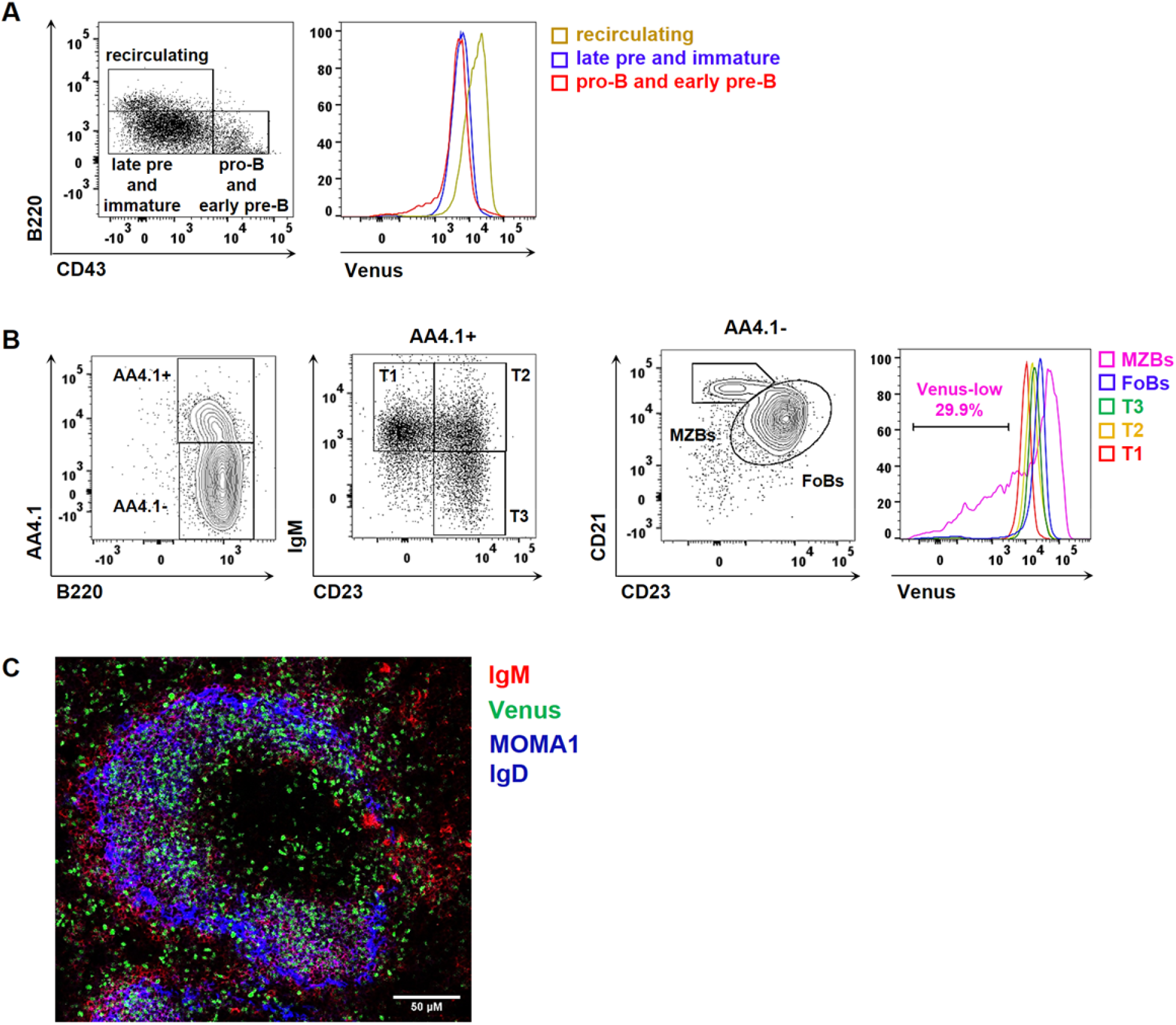
All B cells are recipients of a Notch signal. **(A)** Representative FACS analysis of B220^+^ lymphocytes in the BM of CBF:H2B-Venus mice. B220^+^ cells were subdivided in pro- and early pre-(CD43^+^B220^low^) B cells, late pre- and immature (CD43^low^B220^low^) B cells and recirculating (CD43^-^B220^high^) B cells. n=5. **(B)** Splenic transitional B cells were gated as AA4.1^+^B220^+^ and subdivided into T1-T3 cells according to their differing CD23/IgM expression. Mature B cells were gated as AA4.1^-^B220^+^ and further separated into FoB (CD23^+^CD21^low^) and MZB cells (CD23^low^CD21^high^). n=5. **(A-B)** The histogram overlays show the Venus expression in the previously gated cell populations. The gate depicts the frequencies of Venus-low MZB cells (pink). n=5. **(C)** Immunofluorescence analysis of splenic sections, stained for MZB cells (IgM^+^, red), FoB cells (IgD^+^; blue) and metallophilic macrophages (MOMA1^+^; blue). Green dots represent Venus-expressing cells. The scale bar represents 50 μM. Image is representative for n=3 mice.

In the spleen, Venus expression gradually increased from T1 over T2 to T3 cells, and continued to increase in FoB and MZB cells (Fig. 1B). Although Venus expression was the highest in MZB cells, around one-third of them expressed less Venus than the majority of B cells in the spleen, suggesting that some MZB cells downregulate or turn off Notch signaling over time (Fig. 1B). This observation was recapitulated by our immunohistochemical staining, which showed that some MZB cells are indeed Venus^-^ (Fig. 1C).

### Notch2 signaling is downregulated in most GCB cells and PCs

To analyze the activity of Notch2 signaling in B cell populations that specifically arise during the primary TD immune response, we injected CBF:H2B-Venus mice with 4-Hydroxy-3-nitrophenylacetyl-Chicken Gamma Globulin (NP-CGG) and subsequently analyzed them 7- and 14-days post immunization (p.i.). The Venus signal was progressively downregulated in most germinal center B (GCB) cells, compared to non-GCB cells (Fig. 2A). Furthermore, we found that only a small number of GCB cells were still Venus^high^, with a higher percentage of Venus^high^ cells at day 7 than at day 14 p.i. (Supplementary Fig. 1A). Also, in immunohistochemistry, the GC structures appeared mostly Venus^-^, with only a few Venus^high^ GL7^+^ cells detected. (Fig. 2B). By day 14, the Venus^high^ GCB cells were substantially enriched in the light zone (LZ), whereas Venus^mid^ and Venus^neg^ cells were preferentially found in the dark zone (DZ) (Supplementary Fig. 1B). Furthermore, we found that the Venus expression was strongly reduced in most TACI^+^CD138^+^ PCs (Fig. 2C) and histology for Irf4-expressing cells confirmed this finding (Fig. 2D). Although most PCs had completely shut off the Venus signal, a small portion of TACI^+^CD138^+^ cells remained Venus^high^. Approximately half of these cells still expressed CD19 and B220, suggesting they were more likely (pre-) plasmablasts. On the other hand, the Venus^mid+low^ PCs were uniformly CD19^-^ B220^-^, indicating they are fully mature PCs (Fig. 2C). Taken together, these data imply that Notch signaling is progressively downregulated during the course of terminal PC differentiation.

**Fig. 2.**
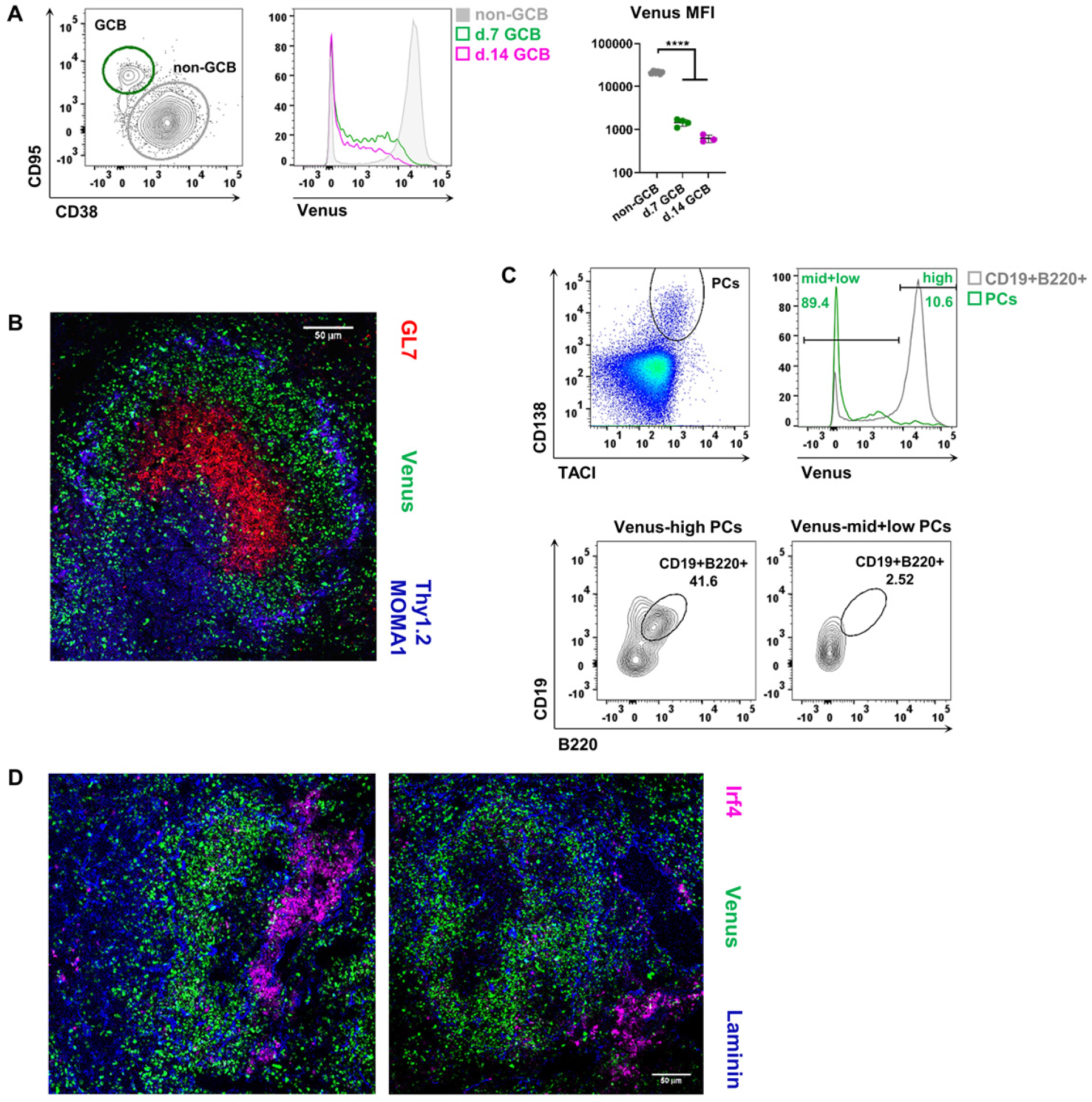
Notch signaling is downregulated in most GCB cells and mature PCs. **(A)** FACS plots are pre-gated on B220^+^ B lymphocytes and show the gating of GCB (CD95^high^CD38^low^) and non-GCB (CD95^low^CD38^+^) cells at day 7 p.i.. The histogram shows an overlay of the Venus expression in GCB cells at day 7 and day 14 p.i. and in non-GCB cells. The graph summarizes median fluorescent intensities (MFI) of Venus in the analyzed populations. ****p<0.0001, Tukey’s one-way ANOVA of logarithmized MFI values. DF=10, F=608.4. n=4 at day 7 p.i., n=3 at day 14 p.i.. **(B)** Splenic sections were stained for GCs (GL7^+^; red), T cells (Thy1.2^+^; blue) and metallophilic macrophages (MOMA1^+^; blue) 7 days p.i. Scale bar represents 50 μm. n=2. **(C)** Splenocytes were pre-gated with a large lymphocyte gate and analyzed for PCs (TACI^+^CD138^+^). The histogram shows an overlay of Venus expression in PCs and B220^+^CD19^+^ B cells. Gates and percentages refer to the green histogram of the PCs, gated as shown in the plot. Contour plots depict the CD19/B220 expression in Venus^high^ and Venus^mid+low^ PCs (TACI^+^CD138^+^). The FACS analyses are representative for day 7 p.i.. n=4. **(D)** Staining of splenic sections to detect plasmablasts and PCs (Irf4^+^; pink) and basement membranes of endothelial cells (Laminin; blue) at 7 days p.i.. Scale bar represents 50 μm. n=2.

### Generation of GCB cells is impaired by Notch2IC induction

Our data thus far revealed that most B cells receive a Notch signal, but Notch signaling is downregulated in the GC. To further study the influence of Notch signaling on the GC reaction, we immunized conditional transgenic mice in combination with the Cγ1-Cre strain (24), in which the Notch2-receptor is deleted (N2KO//CAR) or constitutively activated (N2IC/hCD2) upon TD immunization. As controls, we used control/CAR mice, in which the reporter human coxsackie/adenovirus receptor (CAR) is expressed upon Cre-mediated recombination (25). To trace the B cells that successfully underwent Cre-mediated recombination, we used the reporter gene CAR in both N2KO//CAR and control/CAR mice, and hCD2 in N2IC/hCD2 mice. The mouse strains are described in Materials and Methods and in Supplementary Fig. 2A in more detail.

Following intraperitoneal NP-CGG injection, the deletion efficiency of the targeted regions was comparable in all three mouse strains. The amounts of reporter^+^ cells peaked at day 14 with around 5-6%, followed by a decline to 2% at day 30 (Supplementary Fig. 2B). Although all mice had similar percentages of reporter^+^ B cells, mice with induced Notch2IC-expression were unable to generate GCB cells (Fig. 3A). Thus, only a few GCB cells could be detected in these animals, most of which were reporter^-^ (Fig. 3B).

**Fig. 3.**
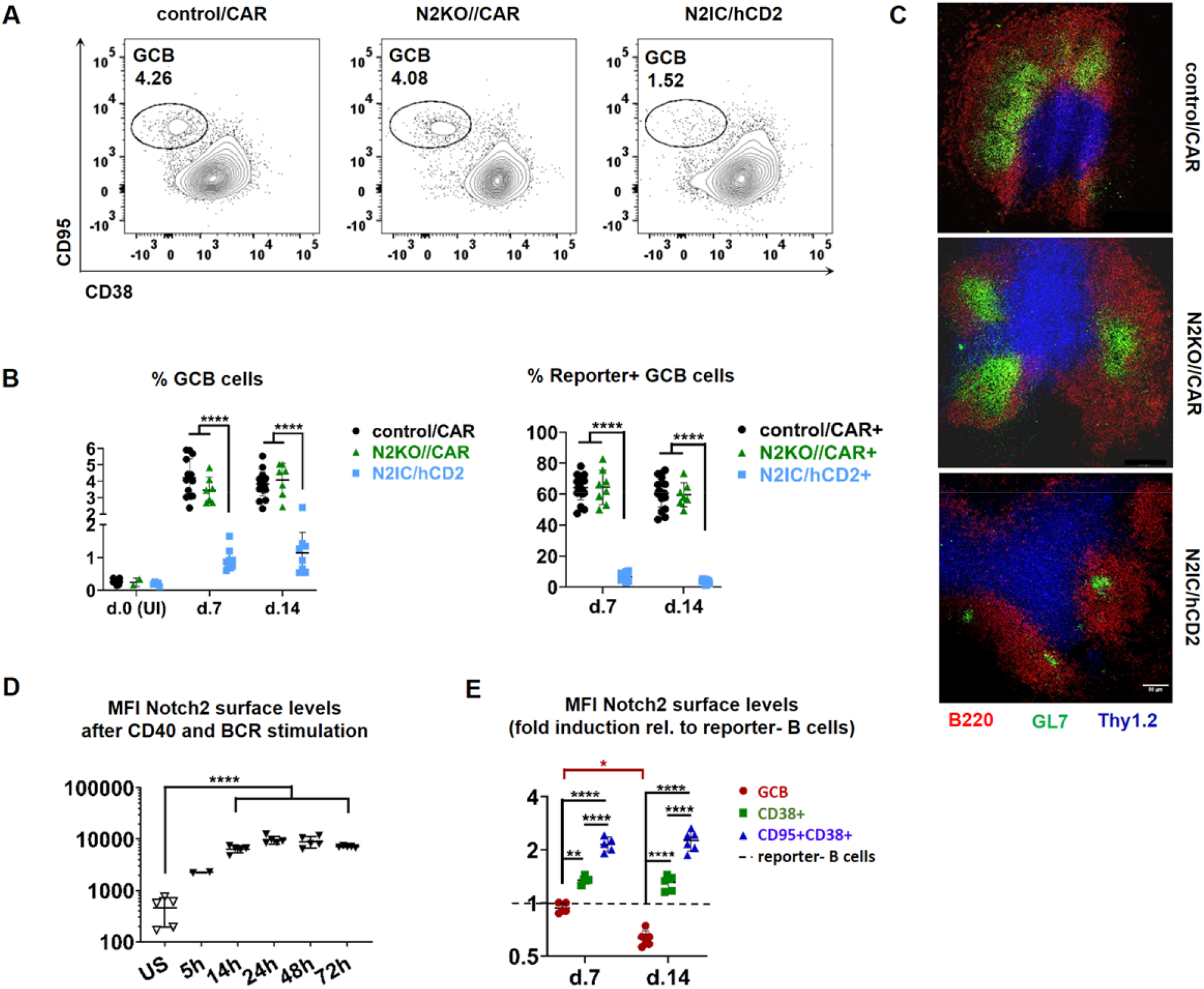
Antigen-induced Notch2IC-expression inhibits GC formation. **(A)** Splenic B220^+^ B cells were analyzed for GCB cells (CD95^+^CD38^low^). FACS plots are pre-gated on B lymphocytes and are representative for day 7 p.i. and for n=14 control/CAR, n=7 N2KO//CAR and n=9 N2IC/hCD2 mice. **(B)** The graph on the left compiles the percentages of total GCB cells in the spleen of mice from the indicated genotypes at the indicated time points p.i.. The graph on the right shows a summary of the percentages of reporter” GCB cells. d0 n=10/2/5, d7 n=14/7/9, d14 n=20/7/8 for control/CAR/N2KO//CAR/N2IC/hCD2 mice, respectively, ****p≤0.0001, ordinary 2-way-ANOVA, Tukey’s multiple comparison test. **(C)** Splenic sections from mice of the indicated genotypes 7 days p.i.. Sections were stained for GCB cells (GL7^+^; green), B cells (B220^+^; red) and T cells (Thy1.2^+^; blue). Scale bar represents 50 μm. n=4 mice per genotype. **(D)** Graphical summary of median fluorescent intensities (MFI) of Notch2 cell surface expression on purified FoB cells at indicated time points after combined α-IgM and α-CD40 stimulation. n=5, except for 5h where n=2. ****p ≤ 0.0001, Tukey’s 1-way-ANOVA. DF=21, F=1.784. Representative histogram overlays of Notch2 surface expression over time are given in Supplementary Fig. 3A. **(E)** Fold induction of Notch2 cell surface expression (MFI) in the reporter” CD38/CD95 subpopulations from control/CAR mice, relative to its expression levels (MFI) in reporter^-^ (CAR^-^) B cells (=1, depicted by the dotted line). d7 n=5, d14 n=6. *p=0.011, **p=0.001 and ****p≤0.0001, Sidak’s 2-way-ANOVA. Representative FACS analysis of the gating strategy is depicted in Supplementary Fig. 3B.

In addition, immunohistochemistry revealed only small GC structures, which were not localized at the T-/B-cell border in N2IC/hCD2 mice (Fig. 3C). In contrast, N2KO//CAR and control/CAR mice generated similar percentages of GCB cells, which clustered into structures with correct localization inside the splenic follicle at the B/T cell border (Fig. 3A-C). These data suggest that Notch2 signaling is not necessary for GCB cell differentiation, and its activation even interferes with the formation of germinal centers.

### Notch2 surface expression is upregulated on activated FoB cells, but downregulated on GCB cells

Previous studies revealed that Notch2 signaling is inversely correlated with GC formation (15, 26). These findings are in line with our observations that N2IC/hCD2 mice are almost completely devoid of GCB cells and Notch signaling is progressively downregulated in GCB cells of CBF:H2B-Venus mice. On the other hand, it has been shown that Notch2 receptor surface expression is upregulated upon BCR stimulation (14) and that Notch2 signaling strongly synergizes with BCR and CD40 signaling in FoB cells, in order to enhance and sustain prolonged B cell activation *in vitro* (27). Therefore, we asked whether in the course of TD immune responses, CD40+BCR stimulation initially results in the upregulation of Notch2 surface expression, which is then downregulated again in GCB cells. We found an around 20-fold upregulation of Notch2 on the cell surface of FoB cells upon combined CD40- and BCR-stimulation *in vitro*. This upregulation started early and peaked between 24h and 48h of stimulation (Fig. 3D, Supplementary Fig. 3A). *In vivo*, Notch2-receptor-expression was approximately 2-fold upregulated in reporter^+^CD38^+^CD95^+^ non-GCB cells in comparison to reporter^-^ B cells, while CD38^-^CD95^+^ GCB cells progressively downregulated Notch2-surface expression over time post immunization (Fig. 3E, Supplementary Fig. 3B). Thus, our *in vitro* and *in vivo* data suggested that Notch2 surface expression is indeed induced on activated non-GCB cells, whereas it is downregulated in fully differentiated GCB cells, consistent with the low levels of Venus expression in GCB cells from immunized CBF:H2B-Venus mice.

### Notch2-signaling in activated FoB cells induces Irf4 expression and inhibits Bcl6 upregulation

Although N2IC/hCD2 mice did not form GCs, they had similar frequencies of reporter^+^ B cells as control/CAR animals (Supplementary Fig. 2B). Most of them expressed high CD38 levels, but varied in their expression of CD95 (Fig. 4A–B). Intracellular staining for the transcription factors Irf4 and Bcl6, which are key regulators of PC- and GCB cell-differentiation respectively (28, 29), revealed an inverse regulation in N2IC/hCD2 and control/CAR mice. Most control/CAR^+^ B cells strongly induced Bcl6 expression, but kept Irf4 levels very low, consistent with their GCB cell phenotype. In contrast, N2IC/hCD2^+^ B cells did not upregulate Bcl6, but showed a strong induction of Irf4 (Fig. 4C). These findings indicate that Notch2 signaling interferes with GC formation through Bcl6 suppression. In addition, the induction of Irf4 expression led us to suspect that Notch2 activation in B cells may support their entry into PC differentiation instead.

**Fig. 4.**
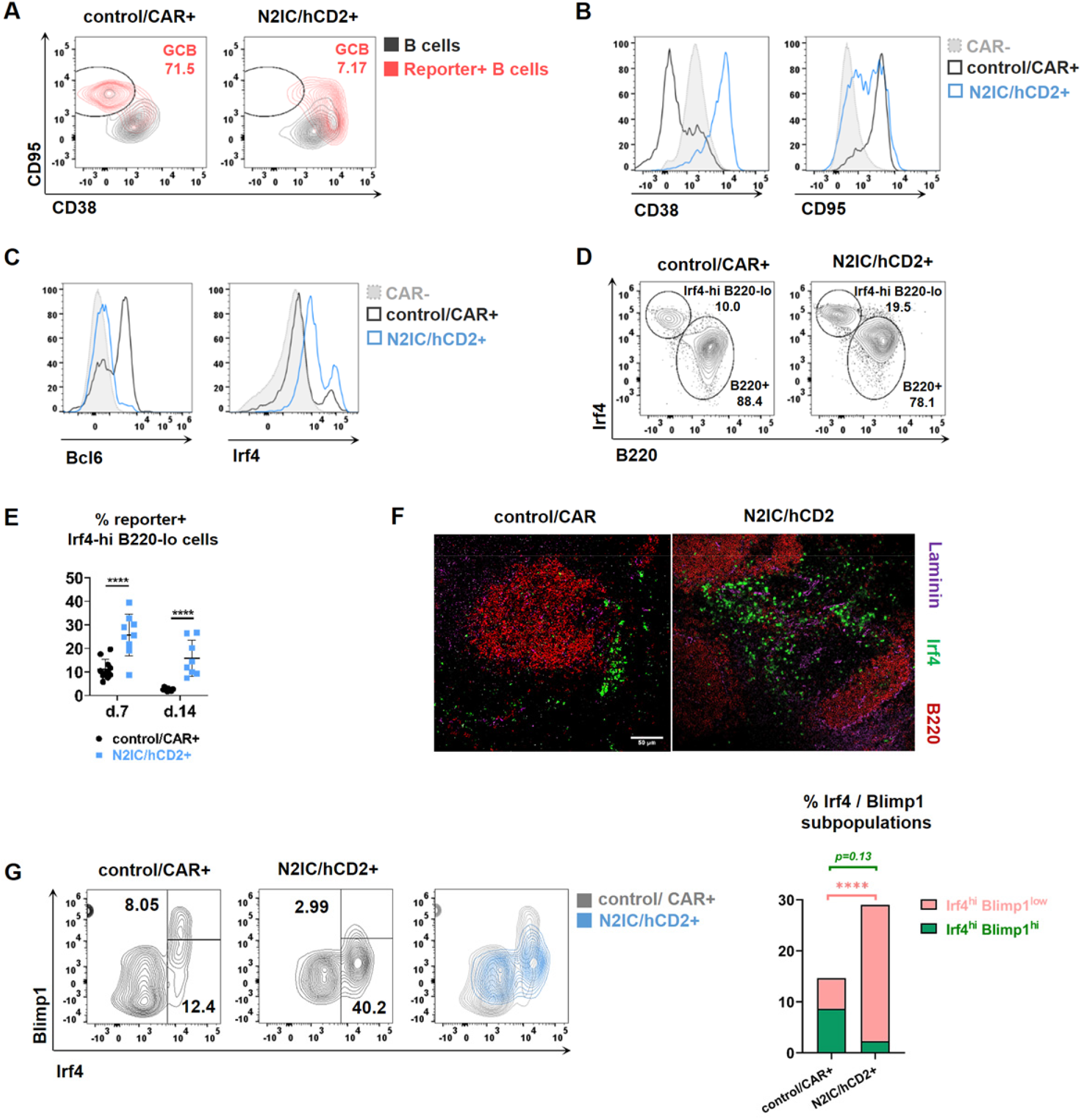
N2IC-expression suppresses Bcl6, but induces Irf4 expression, resulting in enhanced differentiation into plasmablasts. All FACS plots are pre-gated on lymphocytes and are representative for day 7 p.i.. **(A)** Splenic reporter^+^B220^+^ B cells were analyzed for their GCB cell phenotype (CD95^+^CD38^low^). Overlays between reporter^+^ B cells (red) and total B220^+^ cells (black) are shown. Gates and percentages refer to the reporter^+^ populations. **(B)** Histograms depict the overlays of CD38 and CD95 expression in reporter^+^ B cells from N2IC/hCD2 and control/CAR mice, compared to reporter^-^ control B cells. **(A-B)** Analyses are representative for n=14 control/CAR and n=9 N2IC/hCD2 mice. **(C)** Histograms show the overlay of Bcl6 and Irf4 expression in reporter^+^ lymphocytes and reporter^-^ lymphocytes. **(D)** Irf4/B220 expression in reporter^+^ lymphocytes. Two subpopulations were identified: Irf4^high^B220^low^ and B220^+^. **(C-D)** Analyses are representative for n=11 control/CAR and n=9 N2IC/hCD2 mice. **(E)** Summary graph depicting the percentages of reporter^+^Irf4^high^B220^low^ cells at indicated time points. d7 n=11/9, d14 n=11/8 control/CAR mice/N2IC/hCD2 mice, respectively. ****p<0.0001, Sidak’s 2-way-ANOVA. **(F)** Immunofluorescence analysis of splenic sections, stained for (pre-) plasmablasts and plasma cells (Irf4^high^; green), B cells (B220^+^; red) and basement membranes of endothelial cells lining the MZ sinus (Laminin; purple). Scale bar represents 50 μm. n=3 mice per genotype. **(G)** The staining was performed after fixation and permeabilization of cells. Reporter^+^ cells were separated into Blimp1^high^Irf4^high^ mature PCs and Blimp1^low^Irf4^high^ plasmablasts. The contour plot overlay shows the Irf4/Blimp1 expression of control/CAR^+^ and N2IC/hCD2^+^ cells. The stacked bar chart compiles the percentages of Blimp1^high^Irf4^high^ and Blimp1^low^Irf4^high^ cells at day 7 p.i. n=6 control/CAR mice, n=5 N2IC/hCD2 mice. ****p<0.0001, Sidak’s 2-way-ANOVA.

### Notch2 signaling in activated FoB cells enhances plasmablast differentiation

Next, we examined the phenotype of the Irf4-expressing B cells in N2IC/hCD2 mice in more detail. Even though most N2IC/hCD2^+^ cells were Irf4^+^, we found a second fraction of cells, which were Irf4^high^. This fraction was also detected in reporter^+^ cells from control/CAR mice, but at a significantly lower frequency at both day 7 and 14 p.i. compared to N2IC/hCD2 mice. The Irf4^high^ cells were uniformly B220^-^ in both genotypes (Fig. 4D and E), pointing to a plasmablast/PC phenotype. Immunohistochemistry revealed that the enriched Irf4^high^ cells were predominantly located in the extrafollicular regions between neighboring B cell follicles, where plasmablasts and short-lived PCs are usually located (Fig. 4F). Further characterization of the Irf4^high^B220^-^ cells revealed that, in comparison to controls, the expanded Notch2IC-expressing Irf4^high^B220^-^ cells had lower Blimp1 levels, indicating that they were arrested at the plasmablast stage and did not develop into mature PCs (Fig. 4G). This observation is consistent with the data from Notch-reporter mice, which displayed high levels of Venus in pre-plasmablasts and plasmablasts, but low levels in terminally differentiated PCs. ELISpot analyses with splenic B cells revealed increased abundance of NP-specific IgM antibody-secreting cells (ASCs), but severely reduced NP-specific IgG1 and IgG3 cells in N2IC/hCD2 mice related to controls at day 7 p.i. (Supplementary Fig. 4A). A similar trend was visible in the BM, where frequencies of NP-specific IgM^+^ ASCs were increased and the frequencies of IgG1^+^ ASCs were decreased (Supplementary Fig. 4B). In the sera, NP-specific IgM titers were comparable between genotypes, whereas IgG1 titers were decreased in N2IC/hCD2 mice related to control/CAR mice (Supplementary Fig. 4C). In summary, these data suggest that a large proportion of activated FoB cells, which receive a strong Notch signal upon TD immunization, differentiate into extrafollicular plasmablasts, most of which are short-lived (Blimp1^mid^), unswitched, IgM-producing cells.

### Some antigen-activated N2IC-expressing B cells differentiate into MZB cells in the course of the TD immune response

While most reporter^+^ B cells from N2IC/hCD2 mice were Bcl6^-^, they all uniformly expressed higher levels of Irf4 (Irf4^mid^) than the reporter^+^ B cells from control/CAR mice (Fig. 5A–B). Considering the role of induced constitutively active Notch2 signaling in the differentiation of FoB into MZB cells under steady-state conditions (14), we analyzed whether the reporter^+^B220^+^Irf4^mid^Bcl6^-^ cells from N2IC/hCD2 mice adopt a MZB cell phenotype over time. Indeed, at day 7 post-immunization, around one-fourth of N2IC/hCD2^+^B220^+^Irf4^mid^Bcl6^-^ cells had a MZB cell phenotype according to their CD21/CD23 surface expression (Fig. 5C).

**Fig. 5.**
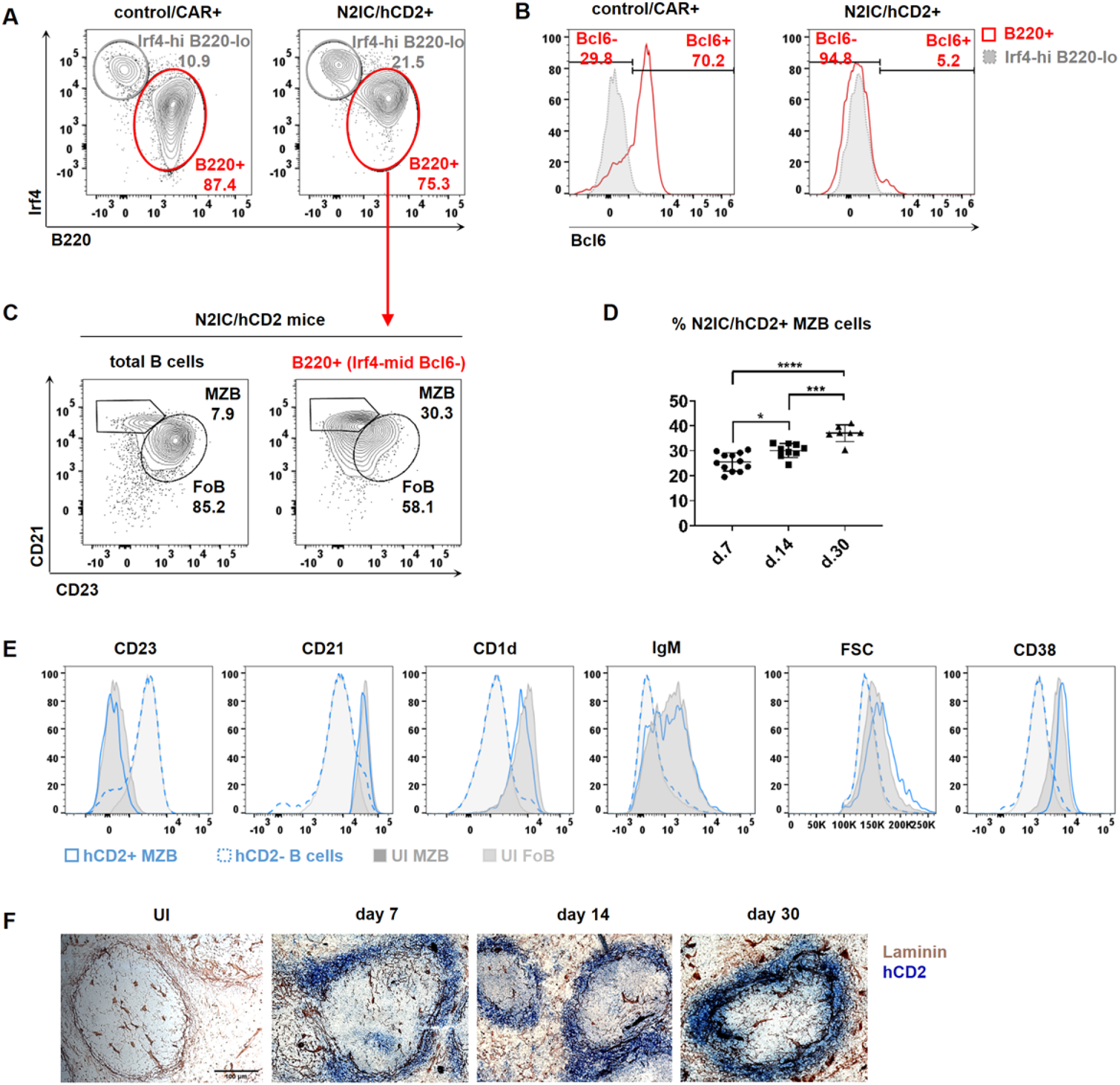
N2IC/hCD2^+^ B cells differentiate into MZB cells with the correct surface phenotype and splenic localization. **(A)** Reporter^+^ lymphocytes were separated into B220^low^Irf4^high^ and B220^+^ cells. **(B)** Histograms show overlays of the Bcl6 expression in reporter^+^B220^+^ and reporter^+^B220^low^Irf4^high^ cells. **(C)** Gating of MZB (CD21^high^CD23^low^) and FoB (CD23^high^CD21^low^) cells in N2IC/hCD2 mice. The left plot is gated on total B220^+^ cells, and the right plot on N2IC/hCD2^+^B220^+^ cells (further described as Irf4^mid^Bcl6^-^ in 5A-B). **(A-C)** FACS plots are representative for day 7 p.i. n=7 control/CAR and n=4 N2IC/hCD2 mice. **(D)** Graphical summary of the percentages of N2IC/hCD2^+^CD21^high^CD23^low^ MZB cells at indicated time points. The gating strategy is shown in Supplementary Fig. 5 (d7 n=12, d14 n=9 and d30 n=7). *p=0.023, ***p=0.0003 and ****p<0.0001, Tukey’s 1-way-ANOVA. DF=25, F=26.71. **(E)** Histogram overlays of the expression of indicated markers within N2IC/hCD2^+^ MZB cells and total splenic hCD2^-^ B cells at day 7 p.i., compared to total MZB and FoB cells from unimmunized (UI) N2IC/hCD2 animals. Analyses are representative for n=9 immunized N2IC/hCD2 mice and n=4 UI N2IC/hCD2 mice. **(F)** Chromogenic immunohistochemical analysis for the localization of hCD2^+^ cells at the indicated time points. Splenic sections were stained for hCD2 (blue) and for basement membranes of endothelial cells lining the MZ sinus with Laminin (brown). Scale bar represents 100 μM. n=3 mice per time point.

Frequencies of these cells increased with time after immunization (Fig. 5D, Supplementary Fig. 5). In parallel, these cells also increased their size (FSC) and upregulated IgM, CD1d and CD38 (Fig. 5E), which is fully in accord with the published MZB cell surface phenotype (CD23^low^CD21^high^IgM^high^CD1d^high^CD38^high^).

Moreover, the N2IC-expressing cells migrated towards the marginal zone. While some N2IC/hCD2^+^ cells were still located within the follicle at day 7, progressively more of these cells were localizing in the MZ with ongoing time post immunization (Fig. 5F). By day 30, most of the N2IC/hCD2-expressing cells were at their defined splenic location, forming an intact MZ ring around the follicle. Together, our data imply that Notch2 signaling in antigen-activated FoB cells not only enhances their entry into PC differentiation, but also induces their differentiation into MZB cells with full cell surface marker phenotype and correct splenic localization in the MZ.

### MZB cells are regularly generated during TD immune responses in a Notch2-dependent manner

We next asked whether MZB cell differentiation occurs only in the transgenic Notch2IC mice or also physiologically in the course of TD immune responses. Indeed, we found reporter^+^CD23^low^CD21^high^ B cells in the spleen of control/CAR mice with unaltered Notch2 signaling (similar to the wildtype situation) after TD immunization (Fig. 6A). The percentages of these cells gradually increased with ongoing time post injection, ranging between 2-10% over the observation period (Fig. 6B).

**Fig. 6.**
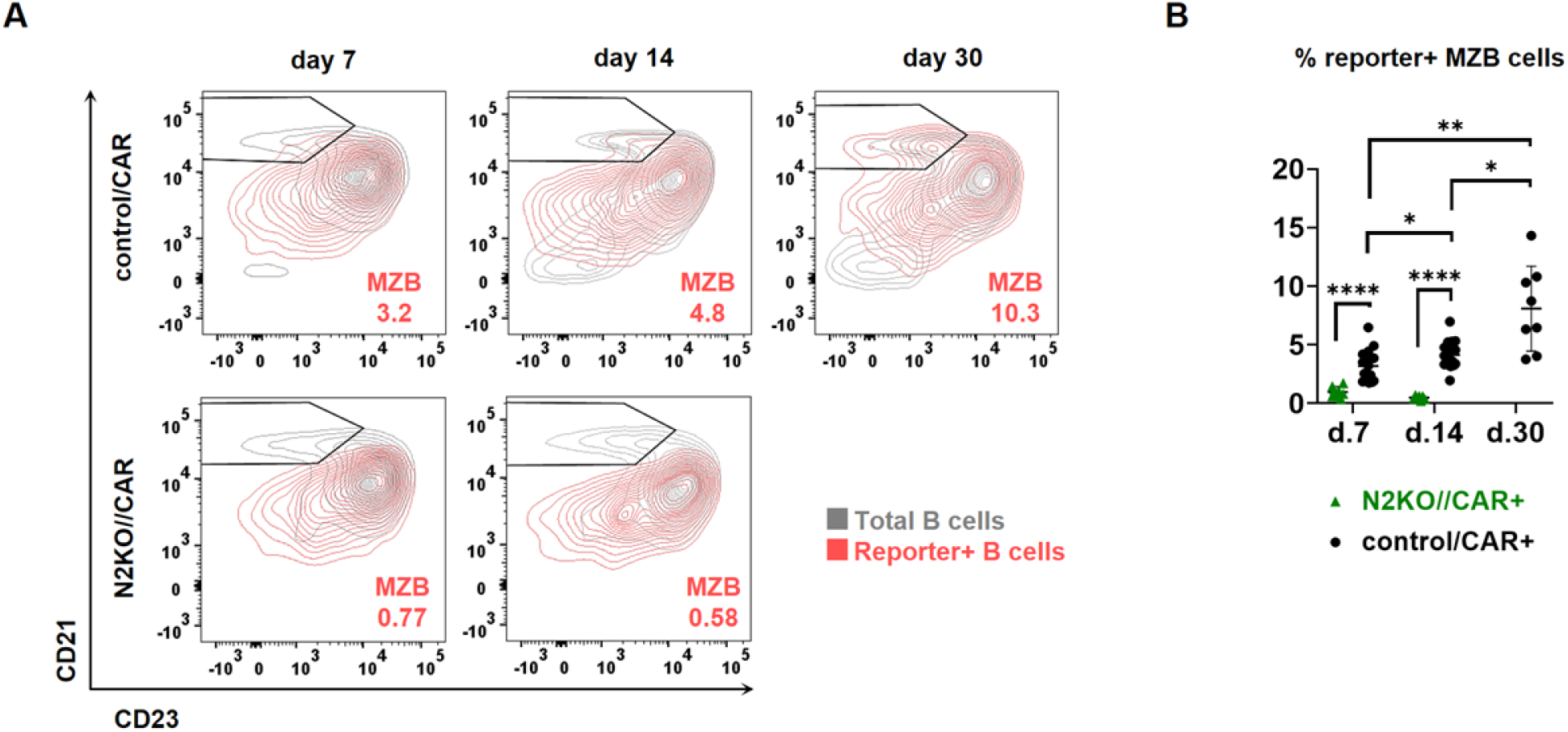
Notch2 signaling is necessary for the generation of reporter_+_ B cells with a MZB phenotype. **(A)** Representative FACS analysis of reporter^+^CD21^high^CD23^low^ MZB cells (red) overlaid with total B220^+^ B cells (grey) in control/CAR and N2KO//CAR mice at the indicated time points. Cells are pre-gated on B220^+^ lymphocytes. Numbers in the FACS plots indicate the percentages of reporter^+^ MZB cells. **(B)** The graph summarizes percentages of reporter^+^ MZB cells at the indicated time points, gated as in (A). Control/CAR mice: d7 n=16, d14 n=17, d30 n=8; N2KO//CAR mice: d7 n=8, d14 n=7. *p=0.002, **p=0.004, ****p<0.0001, Sidak’s 2-way-ANOVA.

Strikingly, reporter^+^CD23^low^CD21^high^ B cells were almost completely abolished in immunized N2KO//CAR mice compared to controls, implying that the generation of MZB cells in the course of the TD immune response is Notch2-dependent (Fig. 6A–B). In addition, the newly generated CAR^+^ MZB cells exhibited high expression of the typical MZB cell surface markers CD1d and IgM, similar to MZB cells in the bulk (Supplementary Fig. 6). Our results established that generation of CAR^+^ MZB cells is a natural outcome of TD immunization and is dependent on the presence of functional Notch2 signaling.

### Quantitatively mapping B cell differentiation during an immune response reveals bifurcation of FoB cells into GCB and MZB cell fates

To gain initial insights about the precursor cells of CAR^+^ MZB cells, we produced correlation plots of CAR-expression kinetics in the FoB, MZB and GCB cell subsets in the course of the TD immune reaction. We employed a sequential gating strategy to determine the frequencies of CAR-expressing B cells among the total GCB cells (CAR^+^B220^+^CD38^low^CD95^high^), FoB cells (CAR^+^B220^+^CD38^+^CD95^mid/low^CD23^+^CD21^low^) and MZB cells (CAR^+^B220^+^CD38^+^CD95^mid/low^CD23^low^CD21^high^). An example of this hierarchical gating is depicted in Supplementary Fig. 7. The correlation analyzes revealed that the rise and decline in CAR^+^ GCB cell numbers mapped linearly with the accumulation of CAR-expressing FoB cells (Fig. 7A), underlining that GCB cells are mainly derived by the activation of FoB cells. We did not find any indication of a precursor-progeny relationship between CAR^+^ MZB and either CAR-expressing FoB or GCB cells, as their CAR-expression kinetics correlated very weakly (Fig. 7B).

**Fig. 7.**
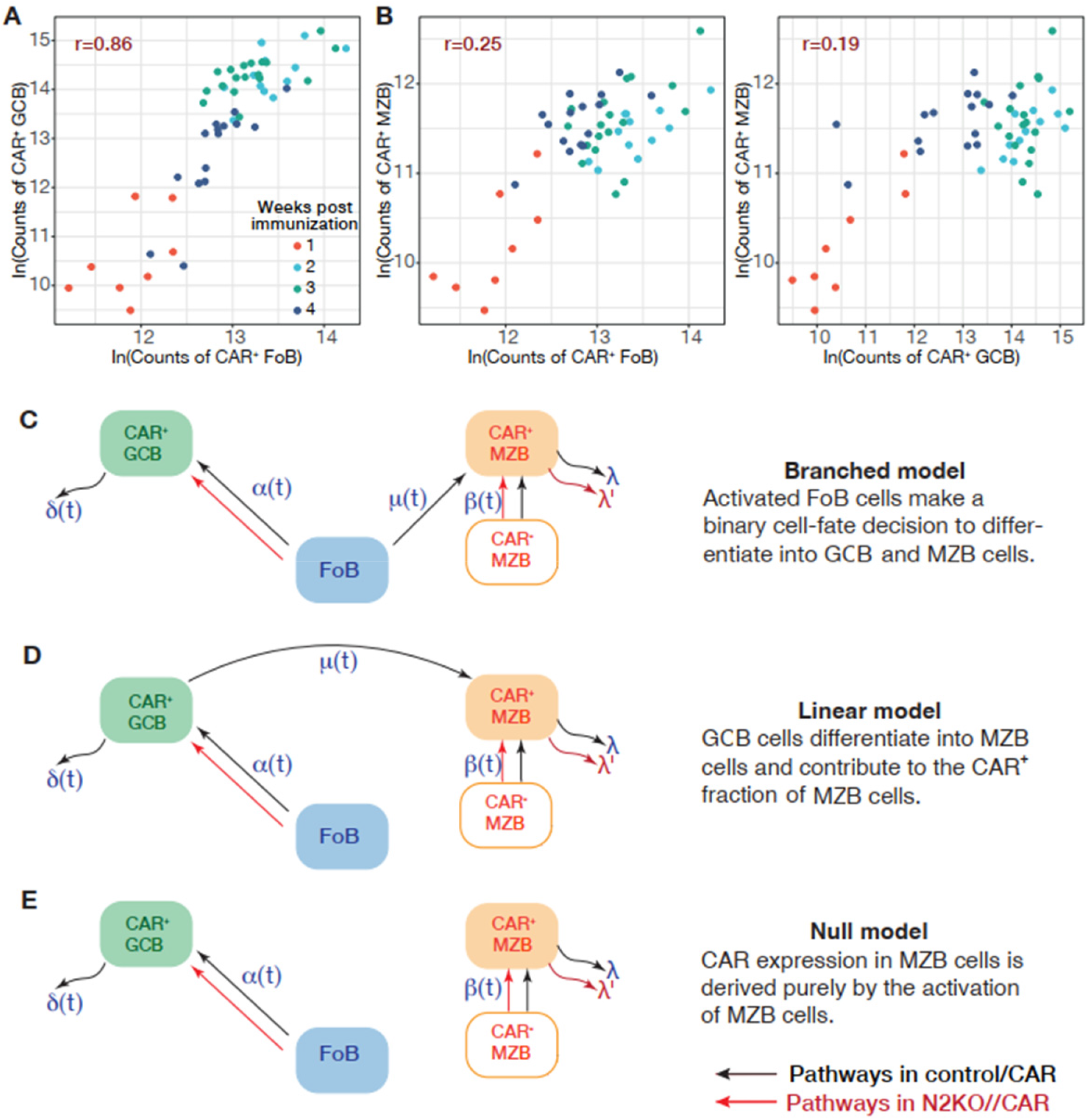
Mathematical modeling of CAR expression dynamics in B cells during an ongoing TD immune response. **(A-B)** Correlation plots of CAR expression kinetics in FoB, MZB and GCB cell subsets. We compare the changes in the logarithmic (ln) counts of CAR^+^ B cells in each subset and show their correlation coefficient ‘*r*’ on each plot. The sequential gating strategy to retrieve the frequencies of each cell population is depicted in Supplementary Fig. 7. The data are grouped in different bins based on the time of observation following TD immunization, indicated by different colors in the plots. d4 n=3, d7 n=9, d9 n=3, d14 n=15, d17 n=3, d22 n=5, d26 n=4, d30 n=6 control/CAR mice after TD immunization. **(C-E)** Schematics of the general forms of the branched, linear and null models of CAR^+^ MZB cell generation upon TD antigen-mediated B cell activation. In-depth details about model specifics are given in the Materials and Methods.

Therefore, to uncover the developmental origins of CAR^+^ MZB cells, we next integrated the data from the immunization experiments into a deterministic mathematical modeling strategy. We hypothesized that CAR^+^ MZB cells emerge either from FoB cells (branched pathway, Fig. 7C) or from GCB cells (linear pathway, Fig. 7D), in addition to potential activation of CAR^-^ MZB cells by the TD antigen, which results in CAR-expression on these MZB cells. We also defined a ‘null model’ in which all CAR^+^ MZB cells are generated only by the activation of CAR^-^ MZB cells (Fig. 7E), without any influx from FoB and GCB cells.

The simplest forms of branched, linear, and null models assume neutral dynamics for the birth-loss of GCB and MZB cells *i.e*., the influx of new cells into the compartment and their net loss (defined as the balance between division and true loss by death and differentiation) remain unchanged over time. To model the effect of antigen availability on B cell dynamics, we considered time-dependent variation in the rate of influx of FoB cells into the GCB cell compartment (*α*(*t*)) and in the rate of upregulation of CAR-expression on MZB cells (*ß*(*t*)). In this ‘time-varying influx’ model, the differentiation of FoB cells into CAR^+^ MZB cells (*μ*) remains unchanged with time (*μ*(*t*) = 0). Additionally, we considered another extension of the neutral model in which the net rate of loss of GCB cells varies with time (δ(*t*)), while the loss rate of CAR^+^ MZB cells (*λ*) remains constant. Model specifics are described in further detail in the Materials and Methods. We fitted each model simultaneously to the time-courses of counts of CAR^+^ MZB cells and CAR^+^ GCB cells from control/CAR mice (red dots in Fig. 8A) and N2KO//CAR mice (blue dots in Fig. 8A), using the Bayesian statistical approach and assessed the relative support for each model using the leave-one-out (LOO) cross validation, calculated as the model weight ‘*W*’ (estimates in Table 1). Details of model weight calculation in Supplementary Text S1. The influxes into the CAR^+^ compartments of GCB and MZB cells were defined using the empirical descriptions of the pool sizes of total FoB cells and CAR^-^ MZB cells (Supplementary Fig. 8 and details in Supplementary Text 2).

**Fig. 8.**
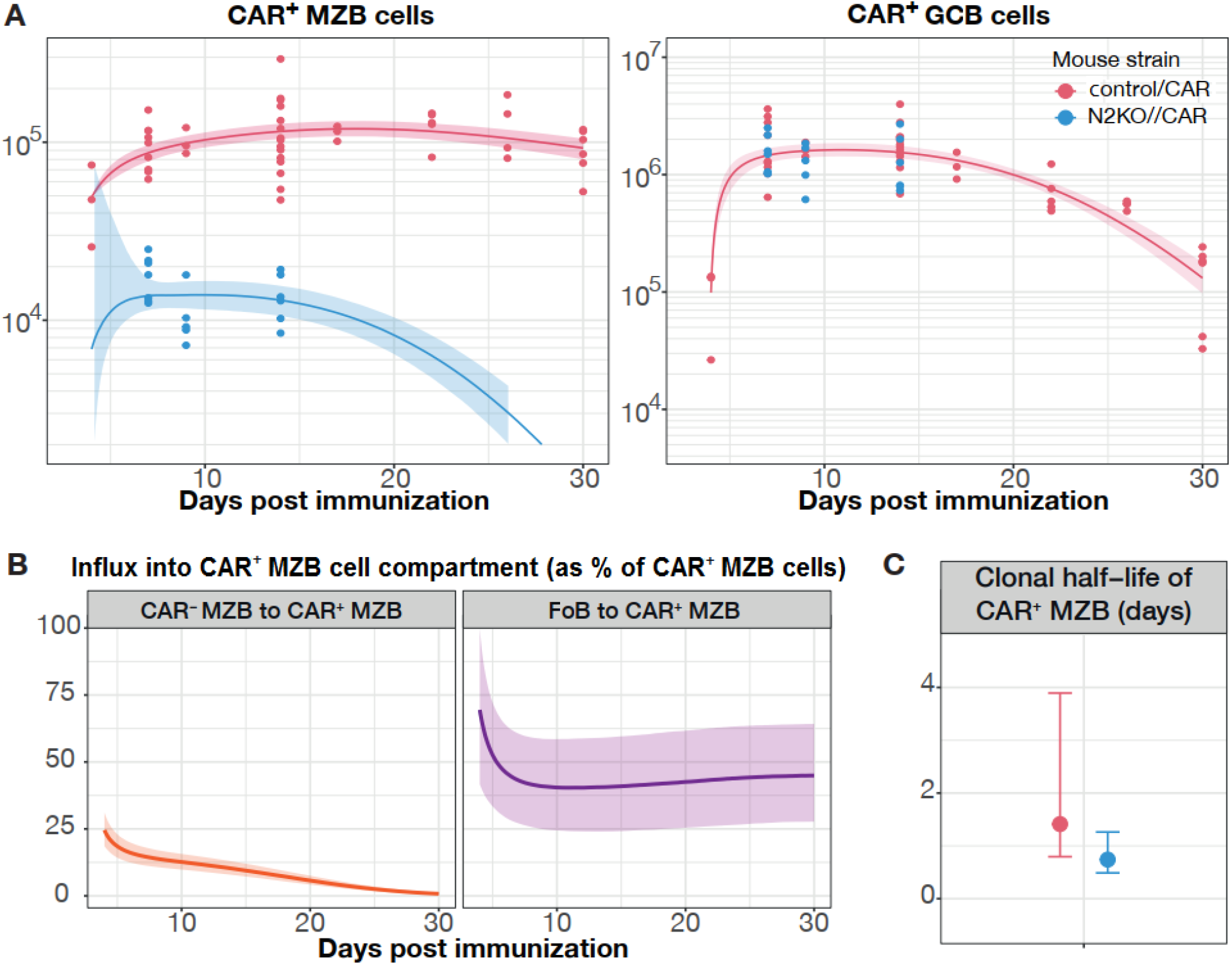
Dynamics of CAR^+^ MZB and GCB cells during a TD immune response. **(A)** Timecourse of counts of CAR^+^ MZB and CAR^+^ GCB cells post immunization in control/CAR (red dots, n=53) and N2KO//CAR (blue dots, n=18) mice. Fits from the best-fitting branched model with time-dependent antigen-activation of FoB and MZB cells (smooth lines) with 95% credible intervals (envelopes). The y-axis depicts total cell counts. The sequential gating strategy to retrieve the frequencies of CAR^+^ GCB and CAR^+^ MZB cells is depicted in Supplementary Fig. 7. Control/CAR mice: d4 n=3, d7 n=9, d9 n=3, d14 n=15, d17 n=3, d22 n=5, d26 n=4, d30 n=6; N2KO//CAR mice: d7 n=7, d9 n=5, d14 n=6. **(B)** The total daily influx of CAR^-^ MZB and CAR^+^ FoB cells into the CAR-expressing MZB cell subset, plotted as percent of total CAR^+^ MZB cell pool sizes (y-axis). **(C)** Clonal half-life of CAR-expressing MZB cells in control/CAR (red) and N2KO//CAR (blue) mice. Clonal half-life is defined as the ln(2)/net loss rate (λ and λ’), which depicts the total time taken for the size of the clonal-lineage to decrease by 2-fold.

**Table 1.**
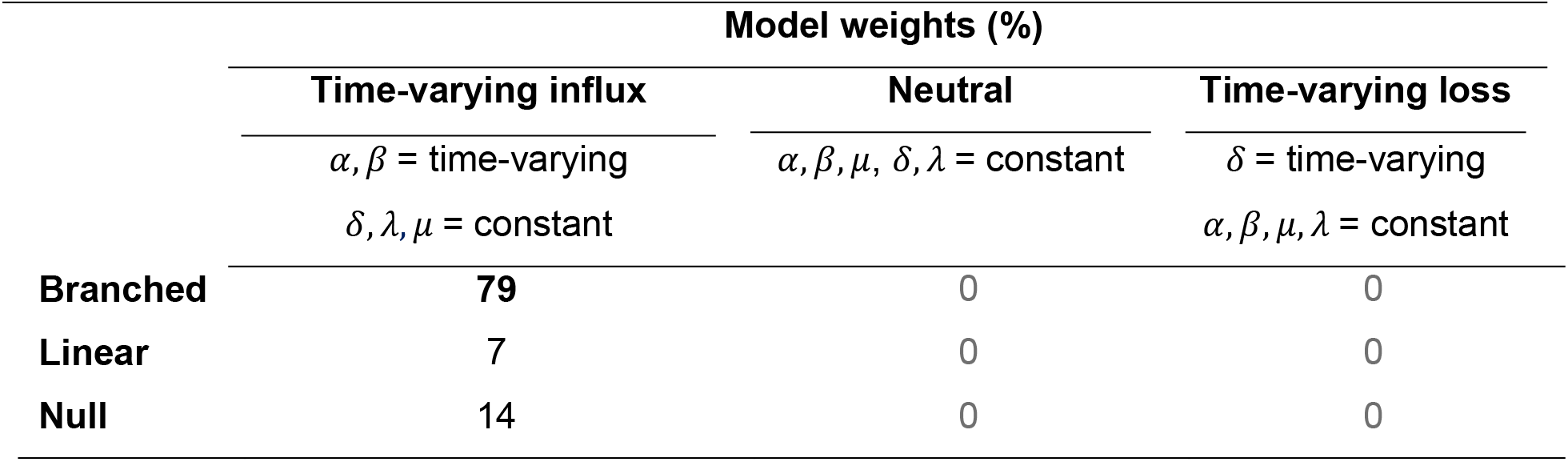
Comparison of models describing population dynamics of MZB and GCB cell populations upon TD immunization. Relative statistical support for each model is indicated as % model weights (see **Supplementary Text 2** for details on model weights estimation). Model specific details are described in Fig. 7C–E; *α* = rate of influx of FoB cells into the GCB cell compartment, *ß* = rate of upregulation of CAR expression on MZB cells, *μ* = rate of differentiation of FoB into CAR^+^ MZB cells, *δ* = loss rate of CAR^+^ GCB cells, *λ* = loss rate of MZB cells. Best-fit model has the highest model weight (depicted in bold).

We found that the combination of the branched pathway with time-varying influx model received the strongest statistical support from the data (*W* = 79%, Table 1). We also explored a version of time-varying influx model in which the rate of differentiation of FoB into CAR^+^ MZB varied with time (*μ*(*t*)), which received poorer statistical support (*W* = 17%), in direct comparison to the best-fitting model (*W* = 79%).

Our best-fitting model nicely captured the rise and fall in CAR^+^ GCB cell numbers and described the persistence of CAR^+^ MZB cells for over a month post immunization (Fig. 8A).

The model estimates that activation of CAR^-^ MZB cells contributes approximately 25% to the CAR^+^ fraction in the MZB cell compartment at day 4. This influx from the CAR^-^ compartment then declines to near-zero by week 3 post immunization (Fig. 8B and Table 2). Differentiation of antigen-activated FoB cells primarily sustained the CAR^+^ fraction in the MZB compartment (Fig. 8B and Table 2), reiterating that Notch2 signaling is required for their development and prolonged maintenance. Consistently, CAR^+^ MZB cell numbers are substantially diminished in Notch2-deficient N2KO//CAR mice (Fig. 8A, blue lines and dots). Our best-fitting model estimated that CAR^+^ MZB cells are lost twice as rapidly in Notch2-deficient mice as compared to the control/CAR mice with functional Notch2 signaling (Fig. 8C and Table 2), indicating that Notch2 signals are also required for maintaining their MZB cell phenotype.

**Table 2.** Parameter estimates from the best-fitting branched pathway with time-varying influx model.

| Parameter | Estimates |  |
| --- | --- | --- |
|  | Median | 95% Credible interval |
| *Per capita* rate of influx of FoB into MZB (day<sup>-1</sup>) | 1/1430 | (1/5500, 1/770) |
| Time taken for the influx rate to halve (days) | 14.2 | (12.9, 15.4) |
| *Per capita* rate of influx of CAR<sup>-</sup> to CAR<sup>+</sup> MZB (day<sup>-1</sup>) | 1/150 | (1/270, 1/100) |
| Rate of net loss of MZB cells in Control/CAR (day<sup>-1</sup>) | 0.49 | (0.18, 0.87) |
| Rate of net loss of MZB cells in N2KO//CAR (day<sup>-1</sup>) | 0.94 | (0.55, 1.4) |

## Discussion

Considering the proximity to the Dll-1-expressing follicular fibroblasts, activation of the Notch2 pathway in most, if not all, B cells in the splenic follicles is a possibility. Thus, it remains unclear, how the differentiation to the MZB cell fate is regulated in B cells receiving a Notch2 signal. We hypothesized that the Notch2 signal-strength determines whether a transitional or FoB cell ultimately develops into a MZB cell. Analyzing the Venus expression in B cells from Notch-reporter mice, which express the yellow fluorescent protein in cells with actively ongoing or recently experienced Notch signals (23) revealed that B cells encounter Notch-ligand-expressing cells in the BM, as well as in the periphery. Venus expression gradually increased throughout stages of B cell development, reaching its highest level in MZB cells, supporting our view that strong Notch2 signaling induces MZB cell development. Surprisingly, around 30% of MZB cells downregulated or completely lost the Venus signal, suggesting that MZB cells do not receive a *de novo* Notch signal in the MZ. By loss of the otherwise high Notch signal, the Venus-low MZB cells may also downregulate their Sphingosine-1-phosphate-receptors (S1PR1 and S1PR3) and migrate back into the follicles (14, 30). Indeed, a high exchange of MZB cells between the MZ and follicular B cell zone has been previously reported, with around 20% of MZB cells migrating from the MZ into the follicle within 1 hour (1). Moreover, inhibition of Notch signaling has been shown to result in the disappearance of MZB cells (31). Thus, oscillation of the Notch2 signal in MZB cells may regulate their migration between the MZ and the follicular region and also, possibly, their expression of MZB cell specific surface markers.

Our data further revealed that Notch2-receptor expression is upregulated on activated FoB cells, whereas it is progressively downregulated on GCB cells in the course of the TD immune response. Fittingly, Venus expression was also strongly and progressively downregulated in GCB cells of CBF:H2B-Venus mice, indicating that Notch signaling is switched off during stages of the GC reaction. Only a small portion of GCB cells, which was enriched in the LZ, was still Venus-high. These cells may receive a Notch signal during the positive selection, when they come into close contact with ligand-providing T follicular helper (T_fh_) cells and follicular dendritic cells (FDCs) (5, 32), the latter having been shown to express high levels of Notch ligands Dll-1 and Jagged-1 (33, 34).

Immunization data from N2IC/hCD2 mice disclosed that B cells receiving a strong Notch signal upon TD immunization adopted a CD38^high^Bcl6^-^Irf4^+^ non-GC phenotype. Within this population, cells which reach very high levels of Irf4 could initiate PC differentiation. Indeed, a larger proportion of Irf4^high^B220^low^ cells was detected in N2IC/hCD2 compared to control/CAR mice, indicating an enhanced PC differentiation upon Notch2 signaling. It has been suggested earlier that Notch1 positively influences the generation of PCs (27, 33). Since Notch2IC-expression most likely mimics both constitutive active Notch1 and Notch2 signaling, it is not excluded that the enhanced PC differentiation observed in N2IC/hCD2 mice is at least in part physiologically caused by Notch1 signaling. However, N2IC-expressing PCs did not reach the same high Blimp1 levels as control PCs, implying that Notch signaling has to be switched off again, to allow for the terminal maturation into PCs. This assumption is supported by the relatively high Venus expression in at least half of the CD19^+^B220^+^ plasmablasts, which is nearly abrogated in almost all of the terminally differentiated CD19^-^B220^-^ PCs in CBF:H2B-Venus (Notch-reporter) mice.

In N2IC/hCD2 mice, a major fraction of reporter^+^B220^+^ B cells adopted a MZB cell surface phenotype and were localized in the MZ, indicating that some B cells that receive a strong Notch-signal in the course of TD immunization and reach Irf4^mid^ levels, differentiate into MZB cells. These reporter^+^ MZB cells were also observed to a lesser extent in control/CAR mice, whereas they were nearly absent in N2KO//CAR mice. These data suggest that MZB cell generation during TD immune responses is a physiological process that is highly dependent on Notch2 signaling, consistent with a recent publication showing that FoB to MZB transdifferentiation is dependent on Notch2 signaling (13). Earlier studies hinted at the antigen-dependent generation of MZB cells (35–39) and clonally related FoB and MZB cells were identified in rats (40). However, it remained unclear whether these MZB cells originate from GC or pre-GC stages. Our mathematical modeling analyses and experimental immunization data favored a Notch2-dependent pathway, in which MZB cells branch out from antigen-activated FoB cells prior to differentiation into the GCB cell stage. Only one-fourth of the CAR^+^ MZB cells seemed to originate from pre-existing (CAR^-^) MZB cells, which responded to the TD antigen activation and therefore upregulated CAR-expression. These cells were generated very early in the TD immune response (day 4), whereas at later time points, the CAR^+^ MZB cell pool was almost entirely sustained by differentiation of antigen-activated non-GC FoB cells. Our best-fit model (branched pathway with time-varying influx) further predicts that CAR-expressing MZB cells have a short clonal half-life (<2 days in control/CAR mice, Fig. 8C). Considering that CAR^+^ MZB cells are generated because they received a strong Notch2 signal in combination with a BCR+CD40 signal, it is likely that they upregulate c-myc and mTOR signaling (41, 42) both of which are described to induce rapid division-independent PC-differentiation of MZB cells (43, 44). This differentiation into PCs may in part be responsible for the rapid loss of CAR^+^ MZB cells. Additionally, MZB cells are known to frequently shuttle back to the follicles, where they most likely replenish their Notch2 signals, as Dll-1 is expressed on follicular fibroblasts (1, 20). Thus, CAR^+^ MZB cells may lose their Notch2-signal in the time course of observation and therefore may migrate back into the follicular B cell zone where they may also lose their typical CAR^+^CD21^high^CD21^low^ MZB phenotype. Of note, our model predicted an even faster loss rate (>2 fold) of reporter^+^ MZB cells in N2KO//CAR mice, compared to controls. We speculate that due to their Notch2-receptor deficiency, the substantially smaller amounts of N2KO//CAR^+^ MZB cells cannot receive a *de novo* Notch2 signal in the follicles. Thus, prolonged Notch2-signal deprivation in these cells likely results in either their complete loss or at least loss of their MZB cell phenotype.

Our modeling data also implied that CAR^+^ MZB cells are largely derived and maintained by antigen-activated FoB cells. This cell pool, described as CD38^+^CD95^mid/low^CD23^+^CD21^low^ B cells, may also contain some memory B cells, especially in the later stages of the GC reaction. Indeed, it has been proposed that GC-derived memory B cells are localized in the follicular B cell zone (45) and are equally distributed between the MZ and the B cell follicles, even 8 weeks after immunization (46). Moreover, memory B cells showed a bimodal expression of CD21 and CD23, suggesting that they have either a FoB or MZB cell phenotype (46). Thus, at least some CAR^+^ MZB cells may originate from memory B cells, which received a strong Notch2 signal either in the GC during positive selection or by DLL-1 expressing fibroblasts in the follicle. In this case, there is a strong indication that the CAR^+^ MZB compartment may be a mixture of memory B cells with somatically mutated Igs on their surface and non-mutated, IgM-expressing ‘antigen-experienced’ cells. This heterogeneity would be apparent in the BCR repertoire of the CAR-expressing MZB cells generated upon immunization. However, previous data showed that antigen-specific memory-like CD80^+^/CD21^low^ cells are more highly mutated than CD80^+^/CD21^high^ cells (46). These findings further support our modeling data, which estimated that the antigen-specific CAR^+^CD21^high^ MZB cells are mainly generated from antigen-activated pre-GCB cells. Nonetheless, understanding the logistics of this dispersion of memory between MZB and FoB cell compartments warrants further investigation, specifically analyzing differentiation trajectories within antigen-experienced B cell lineages.

Overall findings from this study suggest that the human and murine MZB cell compartment may be more similar than previously suspected. The human MZB cell compartment is heterogeneous and its clonal composition appears to change with age (47, 48). In children under the age of two, the majority of MZB cells carry unique BCRs with low mutation frequencies. In contrast, in older children and adults, MZB cells display clonal relationships and are hypermutated, suggesting that naïve MZB cells are progressively replaced by antigen-experienced cells (47). It will be interesting to explore whether the clonal composition and frequencies of hypermutated MZB cells change during aging of mice and whether the ratio of naïve and antigen-experienced MZB cells can be shifted in favor of antigen-experienced MZB cells by frequent antigenic exposures.

## Materials and Methods

### Antibodies used in this study

All details about the different antibodies used in FACS, ELISA/ELISpot and histology are listed in Supplementary Table 1.

### Mouse models

CBF:H2B-Venus reporter mice (23) were purchased from The Jackson Laboratory, backcrossed into the Balb/c background and further maintained in house. Previously described R26/CAG-CARΔ1^StopF^ (25) and N2IC^flSTOP^ mice (15) were crossbred to the Cγ1-Cre strain (24) to generate R26/CAG-CARΔ1^StopF^//γ1-Cre (control/CAR) and N2IC^flSTOP^//γ1-Cre mice (N2IC/hCD2), respectively. R26/CAG-CARΔ1^StopF^ mice carry the human coxsackie/adenovirus receptor CAR in the *rosa26* locus, preceded by a loxP-flanked STOP cassette. Cre recombinase activity results in an expression of a truncated version of the CAR reporter on the cell surface. In the N2IC^flSTOP^ mice, the conditional *notch2IC* allele is directly coupled to the coding sequence for the human CD2 (hCD2) receptor through an IRES site, preceded by a loxP-site-flanked STOP cassette in the *rosa26* locus. Cre recombinase-mediated excision ensures expression of both Notch2IC inside the cell and hCD2 on the cell surface. Lastly, Notch2^fl/fl^ mice (18) were mated to the R26/CAG-CARΔ1^StopF^//γ1-Cre animals to obtain the Notch2^fl/fl^//R26/CAG-CARΔ1^StopF^//Cγ1-Cre (N2KO//CAR) mice. Upon Cre-mediated recombination, exons 28 and 29 of the *notch2* gene, which code for the C-terminal part of RAM23 and the nuclear localization signal (NLS), and the STOP cassette upstream of the *car* gene in the *rosa26* locus are excised. Thereby, Cre-recombined B cells express a truncated non-functional Notch2 receptor (because the following exons 30-33 of *notch2* are out of frame and are thus not translated) and the CAR reporter on the cell surface. Throughout the study, control and mutant mice were age-matched and examined in parallel. All mouse strains were on Balb/c background and analyzed at 10 to 18 weeks of age. Animals were maintained in specific pathogen-free conditions. Experiments were performed in compliance with the German Animal Welfare Law and were approved by the Institutional Committee on Animal Experimentation and the Government of Upper Bavaria.

### Mouse immunizations

To trigger TD immune responses, 10-to 18-week-old mice were injected intraperitoneally with 100 μg of alum-precipitated 4-hydroxy-3-nitrophenylacetyl (NP)-chicken-gamma-globulin (CCG) (Biosearch Technologies, Novato, CA) in 200 μl sterile PBS (Gibco). Mice were analyzed at indicated time points after antigen injection. Respective unimmunized (UI) animals were taken down on the day of each experiment and were used as a day 0 time point.

### Cell purification and *in vitro* cultures

Naïve FoB cells of wildtype Balb/c mice were purified from splenic cell suspension using the MZB and FoB Cell Isolation Kit (Miltenyi Biotech) according to the manufacturer’s instructions. Cells were cultured at a density of 5×10^5^ cells/ml in flat-bottom 96-well plates in RPMI-1640 medium (Gibco), supplemented with 10% fetal calf serum (FCS, PAA Cell Culture Company), 1% L-glutamine, 1% non-essential amino acids, 1% sodium pyruvate and 50 μM β-mercaptoethanol. With the exception of FCS, all supplements were purchased from Gibco. An agonistic antibody against CD40 (2.5 μg/ml; eBioscience HM40-3) and an antibody specific for IgM (15 μg/ml; AffiniPure F(ab’)2 goat anti-mouse IgM, μ-chain, Dianova) were used as a combined stimulus. Cells were harvested at 0 h (unstimulated, US), 5 h, 14 h, 24 h, 48 h and 72 h and analyzed by flow cytometry after staining with an anti-Notch2-PE antibody (BioLegend). Dead cells were discriminated from living cells using a fixable live/dead cell staining kit (Invitrogen).

### Histology

Murine splenic tissues were embedded in O.C.T. compound (VWR Chemicals, USA), snap frozen and stored at −20 °C. Tissues were sliced with a cryostat with 7 μm thickness, mounted on glass slides and stored at −80 °C for immunohistochemical (IHC) or immunofluorescence (IF) analysis.

For the chromogenic IHC, splenic sections were fixed with ice-cold acetone for 10 min, washed with PBS and blocked with 1% BSA, 5% goat serum in PBS. Samples were blocked using the Avidin/Biotin blocking kit (Vector), following the manufacturer protocol. Detection of hCD2-expression was done by staining with a biotinylated mouse-anti-hCD2 antibody and a rabbit-anti-Laminin antibody overnight at 4 °C in 1% PBS/BSA. Secondary antibody staining was done for 1 h at room temperature with streptavidin-coupled alkaline phosphatase and peroxidase-coupled anti-rabbit IgG in 1% PBS/BSA. Enzymatic chromogenic development was done with the AEC substrate and Blue AP substrate kits (Vector), following the manufacturer manual. Slides were embedded in Kaiser’s Gelatin (Carl Roth) and analyzed using an Axioscope (Zeiss) microscope with an AxioCam MRc5 digital camera (Carl Zeiss GmbH) (14)

For the IF staining, sections were fixed for 10 min with PBS-diluted 3% PFA (Histofix, Carl Roth), rinsed in PBS and rehydrated for 8 min in PBS + 50 mM NH4Cl. For the visualization of the MZ, T- and B-cell zone, sections were blocked with 1% BSA, 5% rat serum, 5% chicken serum in PBS for 30 min, followed by Avidin/Biotin blocking (Vector) according to the manufacturer’s protocol. Primary antibodies (goat-anti-mouse IgM, anti-IgD-Biotin, and anti-MOMA1-Biotin), secondary antibodies (chicken anti-goat IgG AF647) and Streptavidin-AF594, were incubated for 1 h at room temperature in 0.5% BSA/PBS. To detect plasmablasts and PCs, sections were permeabilized and blocked for 20 min with 0.3% Triton X, 1% BSA, 5% goat serum in PBS. Primary (rat-anti-mouse Irf4, rabbit-anti-mouse Laminin) and secondary antibodies (goat-anti-rat AlexaFluor488 or goat-anti-rat AlexaFluor647, goat-anti-rabbit Cy3) were incubated in 1% BSA/PBS for 1 h at room temperature. Fluorophore-coupled antibodies (anti-mouse B220-APC) were incubated for 2 h at room temperature, where appropriate. For the detection of germinal center (GC) structures, sections were blocked with 1% BSA, 5% rat serum in PBS for 30 min. Rat-anti-mouse GL7-FITC or GL7-APC antibody was incubated overnight at 4 °C. Directly-coupled antibodies (Thy1.2-Biotin, MOMA1-Biotin, and B220-APC) and Streptavidin-AlexaFluor594 were incubated for 1 h at room temperature in 1% BSA/PBS. Slides were embedded in ProLong Glass Antifade (Invitrogen). Images were acquired on a TCS SP5 II confocal microscope (Leica) and picture stacks were composed in ImageJ (14, 49).

### ELISA

NP-specific antibody titers in the sera were measured by ELISA, following the procedure described by (49, 50). NUNC plates (Nunc) were coated with NP3-BSA or NP14-BSA (10 mg/ml, Biosearch Technologies) in carbonate buffer (0.1 M NaHCO3, pH=9.5) and incubated over night at 4 °C. Plates were washed with PBS, followed by blocking for 2 h with blocking buffer (1% milk powder in PBS for IgG1 ELISA, 5% milk powder in PBS for IgM ELISA). Serum samples were distributed in a 1:2 serial dilution in blocking buffer over 8 wells, starting with 100 μl/well of either 1:10 (IgM) or 1:100 (IgG1) diluted serum in the top row of the plate. The sera were incubated for 1 h at room temperature, followed by washing with PBS. For IgG1 ELISA, the biotinylated antibody was diluted in blocking buffer and incubated for 30 min at room temperature. For IgM ELISA, a directly coupled anti-IgM-HRP antibody was diluted in blocking buffer and incubated for 1 h at room temperature. For the IgG1 ELISA, plates were washed with PBS and incubated with streptavidin horseradish peroxidase (HRP) Avidin D (Vector) diluted in blocking buffer. Following three washing steps with PBS and PBS-T, detection of the HRP signal was done with substrate buffer (1 tablet o-Phenylenediamine (Sigma, P-7288) + 35 mL substrate buffer (0.1 M citric acid, 0.1 M Tris (Sigma)) supplemented with 21 μl H_2_O_2_ (Sigma)). The absorbance was determined with a microplate ELISA reader (Photometer Sunrise RC, Tecan) at an optical density (OD) at 405 nm. To correctly quantify the measured titers and compare different independent assays, internal standards consisting of pooled serums from six to eight NP-CGG immunized mice were used. A calibration curve was produced using the absorbance values measured in the serum standards, which was then used as a guideline to calculate the NP-titers in sera of immunized control/CAR and N2IC/hCD2 animals. All NP-specific titers are given as log10 relative units.

### ELISpot

For antigen-specific ELISpot with splenic or BM cells, 96-well ELISpot plates (Millipore) were coated overnight at 4°C with NP3– or NP14–bovine serum albumin (BSA; Biosearch Technologies), diluted in carbonate buffer. Plates were washed with PBS and blocked for 3 h at 37 °C with 10% FCS/B-cell medium (BCM). 5×10^5^ cells were seeded per well and incubated for 24 h at 37 °C. Plates were washed with PBS-T (PBS with 0.025% Tween20). Biotinylated antibodies against the detecting isotype were diluted in 1% BSA/PBS and incubated for 2 h at 37 °C. After washing with PBS-T, streptavidin horseradish peroxidase Avidin D (Vector) in 1% BSA/PBS was added and incubated for 45 min at room temperature. For the development of spots, a developing buffer was prepared as follows: one tablet of each 3,3’-Diaminobenzidin peroxidase-substrate (0.7 mg/ml, Sigma-Aldrich, gold and silver tablets) was dissolved in 5 ml distilled water, the solution was pooled and filtered. 50 μl/well of this buffer was incubated for 8 min (splenocytes) or 12 min (BM cells). The reaction was stopped by washing with distilled water. Spots were visualized and counted with the ImmunoSpot Series 5 UV Analyzer (CTL Europe) (49, 50).

### Flow cytometry (FACS)

Single cell suspensions were prepared from the spleen and BM. Surface staining of lymphocytes with the corresponding antibodies in MACS buffer (Miltenyi) was performed on ice for 25 min. To omit dead cells from the analysis, cells were stained for 20 min on ice with LIVE/DEAD Fixable Blue Dead Cell Stain Kit (Invitrogen) before the surface antibody staining. For intracellular FACS staining, cells were fixed with 2% paraformaldehyde (1:2 PBS-diluted Histofix) for 10 min at room temperature and permeabilized in ice-cold 100% methanol for 10 min on ice. Cells were incubated for 1 h at room temperature with the corresponding antibodies. Dead cells were excluded by staining for 5 min on ice with LIVE/DEAD Fixable Blue Dead Cell Stain Kit (Invitrogen) before fixation. Cytometry analysis was performed on a LSRII FACS Fortessa (BD Biosciences). Results were evaluated using FlowJo (v9 and v10).

### Statistics

All statistical analyses, including testing for distributions and equal variance, calculations of means and SDs, determining p-values, F values and degrees of freedom of 1-way or 2-way ANOVA with multiple comparisons tests were done with GraphPad Prism (versions 8 through 10). P values of less than 0.05 were considered significant. *P < 0.05, **P < 0.01, ***P < 0.001, and ****P < 0.0001. Error bars in all figures define means ± SD. Samples sizes and the chosen statistical analysis test are indicated in each figure legend.

### Mathematical models of B cell dynamics during a TD immune response

To model the emergence of CAR^+^ MZB and GCB cells post immunization in control/CAR (which are rated as WT controls) and N2KO//CAR mice, we developed an array of system of ordinary differential equations (ODEs) that explored diverse pathways of B cell differentiation and logistics of their maintenance post activation. We observe no significant differences in the counts of CAR^+^ MZB and GCB cells up to day 4 post immunization and therefore anchor the initial condition t_0_ = 4 days in all our models. Model schematics are described in Figure 7.

### Mechanisms of maintenance of GCB cells

We assumed that all GCB cells are derived from antigen-activated FoB cells with a *per capita* rate *α*. We explored modeling possibilities in which either the influx of cells into the GC compartment or their loss from the GC pool are varying with time post immunization.

1. **Neutral dynamics.** This is the simplest model, which assumes constant birth-loss. The rates of influx into the GC (*α*) and net loss of cells from the GC (*δ*), remain constant with time. In all our models, the net loss rate *δ* is the balance between production of new GCB cells by cell division and their ‘true loss’ by death and differentiation.
2. **Time-dependent loss of GCB cells.** This model assumes that the net loss rate of GCB cells varies with time post immunization. We explored the sigmoid form of *δ*(*t*), as shown in equation 1 below. The influx of antigen-activated FoB cells into the GC compartment *α* is assumed to be constant in this model. Parameters *v* and *δ*_0_ are unknowns in this model and are estimated from model fits, along with *α*.

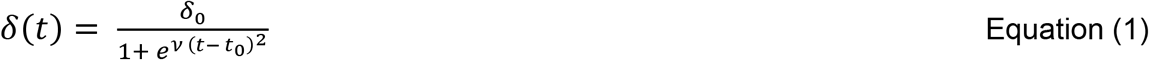
3. **Time-dependent recruitment of FoB cells into GC.** In this model, the *per capita* rate of recruitment of FoB cells into the GC varies with time *α*(*t*) and is defined as,

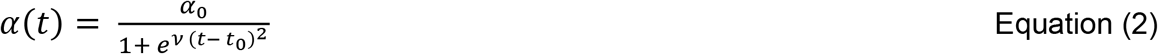 The net loss rate of GCB cells *δ* is constant in this model, estimated from the model fits along with *v* and α_0_.

### Loss of CAR^+^ MZB cells

We assumed a constant *per capita* rate of net loss *λ* for CAR^+^ MZB cells. It has been shown that persistent Notch2 signaling is required for B cells to maintain their MZ status (31). Therefore, the net loss rate of MZB cells reflects the balance between their selfrenewal (division) and their true loss via death, differentiation and deprivation of Notch2-dependent signals. In N2KO//CAR mice, antigen-activation of B cells leads to deletion of Notch2 and thus increases the propensity of their loss by deprivation of Notch2-mediated signals. Therefore, we assume that the net loss rate of CAR^+^ MZB cells in N2KO//CAR mice *λ_N2_* to be higher than their net loss rate in control/CAR mice.

### Mechanisms of generation of MZB cells

We explored three different models – branched, linear and null – to explain the emergence of CAR^+^ MZB cells after TD-immunization.

1. **Branched model:** In this model, the antigen-dependent activation of FoB cells induces a branch point in the B cell differentiation pathway, where some cells participate in GC reactions, while others acquire a MZB cell phenotype (Fig. 7C). Since the dynamics of CAR-expression on FoB cells are unclear, we considered that antigen-activated FoB cells (irrespective of their CAR expression status) differentiate into CAR^+^ MZB cells (M^+^) with a *per capita* rate *μ*. The dynamics for CAR^+^ GCB (G^+^) and CAR^-^ GCB (G^-^) cells are identical between control/CAR and N2KO//CAR mice. The general form ODE system depicting the branched pathway is defined as,

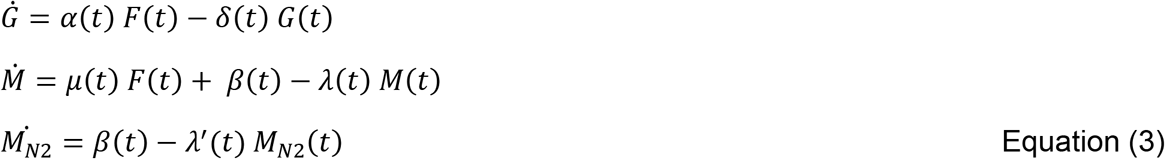
2. **Linear model:** This model follows a linear path in which antigen-activated FoB cells differentiate into GCB cells, which then subsequently give rise to CAR-expressing MZB cells (Fig. 7D). The general form of ODE system of the linear pathway is given as,

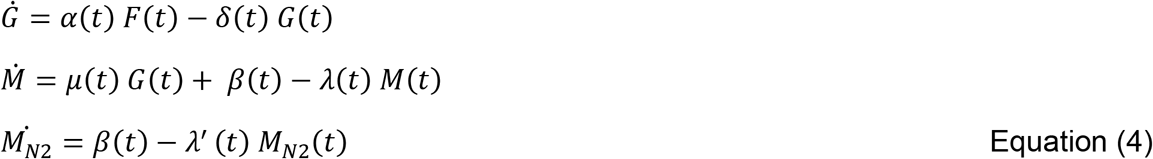
3. **Null model:** This model assumes that the CAR^+^ fraction in MZB cells is purely derived from activation of pre-existing MZB cells and is maintained by activation-induced continuous new deletion of the stop-cassette in CAR^-^ MZB cells and by selfrenewal of newly generated CAR^+^ MZB cells (Fig. 7E).

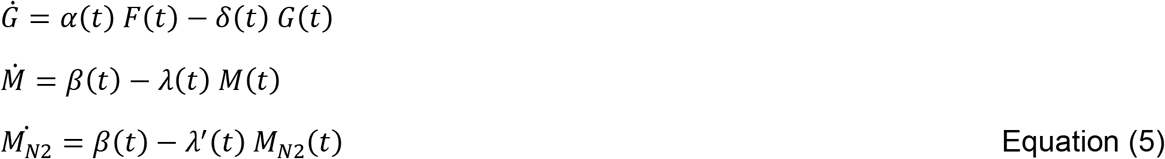 We assumed that the rate of activation of CAR^-^ MZB to CAR^+^ MZB cells *(ß)* varies with time as shown below.

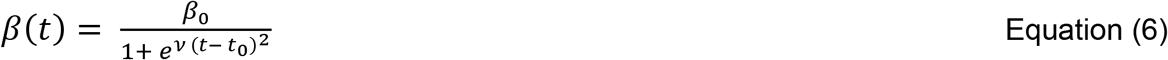 We also explored the possibility of rate of influx of antigen-activated FoB cells to CAR^+^ MZB (*μ*) cells varying with time with the same form as shown in equation 6, likely due to the dependence on antigen availability. Code and data used to perform model fitting are freely available at this linked <u>Github repository</u>.

## Acknowledgements

This work was supported by the Deutsche Forschungsgemeinschaft (DFG ZI1382/4-1). We thank the animal facility of the Helmholtz Center and our animal caretaker team for the excellent housing of the mice and Krisztina Zeller for exquisite technical help with murine DNA preparation and genotyping.

## Authorship Contributions

T.B. performed most experiments and analyzed data. M.L. performed experiments.

A.J.Y. contributed to mathematical modeling and manuscript editing. S.R. developed and performed mathematical modeling. S.R., L.J.S and U.Z.-S designed research and contributed to data interpretation and manuscript preparation. T.B. and U.Z.-S. wrote the manuscript. L.J.S and U.Z.-S. contributed equally. All authors read and approved the final paper.

## Additional Information

Supplementary information: The online version of the submitted paper contains supplementary materials as a single, separate PDF file containing: eight Supplementary Figures (Supplementary Fig. 1-8), one Supplementary Table, Supplementary Notes with two chapters (Supplementary Text 1 and 2) and a list of Supplementary References.

## References

1. Arnon TI, Horton RM, Grigorova IL, Cyster JG. Visualization of splenic marginal zone B-cell shuttling and follicular B-cell egress. Nature. 2013;493(7434):684–8.

2. Martin F, Oliver AM, Kearney JF. Marginal zone and B1 B cells unite in the early response against T-independent blood-borne particulate antigens. Immunity. 2001;14(5):617–29.

3. Cerutti A, Cols M, Puga I. Marginal zone B cells: virtues of innate-like antibody-producing lymphocytes. Nat Rev Immunol. 2013;13(2):118–32.

4. Shlomchik MJ, Luo W, Weisel F. Linking signaling and selection in the germinal center. Immunol Rev. 2019;288(1):49–63.

5. Victora GD, Nussenzweig MC. Germinal Centers. Annu Rev Immunol. 2022;40:413–42.

6. Pillai S, Cariappa A. The follicular versus marginal zone B lymphocyte cell fate decision. Nat Rev Immunol. 2009;9(11):767–77.

7. Allman D, Lindsley RC, DeMuth W, Rudd K, Shinton SA, Hardy RR. Resolution of three nonproliferative immature splenic B cell subsets reveals multiple selection points during peripheral B cell maturation. J Immunol. 2001;167(12):6834–40.

8. Martin F, Kearney JF. Positive selection from newly formed to marginal zone B cells depends on the rate of clonal production, CD19, and btk. Immunity. 2000;12(1):39–49.

9. Srivastava B, Quinn WJ, 3rd, Hazard K, Erikson J, Allman D. Characterization of marginal zone B cell precursors. J Exp Med. 2005;202(9):1225–34.

10. Vinuesa CG, Sze DM, Cook MC, Toellner KM, Klaus GG, Ball J, et al. Recirculating and germinal center B cells differentiate into cells responsive to polysaccharide antigens. Eur J Immunol. 2003;33(2):297–305.

11. Agenes F, Freitas AA. Transfer of small resting B cells into immunodeficient hosts results in the selection of a self-renewing activated B cell population. J Exp Med. 1999;189(2):319–30.

12. Cariappa A, Boboila C, Moran ST, Liu H, Shi HN, Pillai S. The recirculating B cell pool contains two functionally distinct, long-lived, posttransitional, follicular B cell populations. J Immunol. 2007;179(4):2270–81.

13. Gomez Atria D, Gaudette BT, Londregan J, Kelly S, Perkey E, Allman A, et al. Stromal Notch ligands foster lymphopenia-driven functional plasticity and homeostatic proliferation of naive B cells. J Clin Invest. 2022.

14. Lechner M, Engleitner T, Babushku T, Schmidt-Supprian M, Rad R, Strobl LJ, et al. Notch2-mediated plasticity between marginal zone and follicular B cells. Nat Commun. 2021;12(1):1111.

15. Hampel F, Ehrenberg S, Hojer C, Draeseke A, Marschall-Schroter G, Kuhn R, et al. CD19-independent instruction of murine marginal zone B-cell development by constitutive Notch2 signaling. Blood. 2011;118(24):6321–31.

16. Saito T, Chiba S, Ichikawa M, Kunisato A, Asai T, Shimizu K, et al. Notch2 is preferentially expressed in mature B cells and indispensable for marginal zone B lineage development. Immunity. 2003;18(5):675–85.

17. Hozumi K, Negishi N, Suzuki D, Abe N, Sotomaru Y, Tamaoki N, et al. Delta-like 1 is necessary for the generation of marginal zone B cells but not T cells in vivo. Nat Immunol. 2004;5(6):638–44.

18. Besseyrias V, Fiorini E, Strobl LJ, Zimber-Strobl U, Dumortier A, Koch U, et al. Hierarchy of Notch-Delta interactions promoting T cell lineage commitment and maturation. J Exp Med. 2007;204(2):331–43.

19. Tan JB, Xu K, Cretegny K, Visan I, Yuan JS, Egan SE, et al. Lunatic and manic fringe cooperatively enhance marginal zone B cell precursor competition for delta-like 1 in splenic endothelial niches. Immunity. 2009;30(2):254–63.

20. Fasnacht N, Huang HY, Koch U, Favre S, Auderset F, Chai Q, et al. Specific fibroblastic niches in secondary lymphoid organs orchestrate distinct Notch-regulated immune responses. J Exp Med. 2014;211(11):2265–79.

21. Lopes-Carvalho T, Kearney JF. Development and selection of marginal zone B cells. Immunol Rev. 2004;197:192–205.

22. Hammad H, Vanderkerken M, Pouliot P, Deswarte K, Toussaint W, Vergote K, et al. Transitional B cells commit to marginal zone B cell fate by Taok3-mediated surface expression of ADAM10. Nat Immunol. 2017;18(3):313–20.

23. Nowotschin S, Xenopoulos P, Schrode N, Hadjantonakis AK. A bright single-cell resolution live imaging reporter of Notch signaling in the mouse. BMC Dev Biol. 2013;13:15.

24. Casola S, Cattoretti G, Uyttersprot N, Koralov SB, Seagal J, Hao Z, et al. Tracking germinal center B cells expressing germ-line immunoglobulin gamma1 transcripts by conditional gene targeting. Proc Natl Acad Sci U S A. 2006;103(19):7396–401.

25. Heger K, Kober M, Riess D, Drees C, de Vries I, Bertossi A, et al. A novel Cre recombinase reporter mouse strain facilitates selective and efficient infection of primary immune cells with adenoviral vectors. Eur J Immunol. 2015;45(6):1614–20.

26. Valls E, Lobry C, Geng H, Wang L, Cardenas M, Rivas M, et al. BCL6 Antagonizes NOTCH2 to Maintain Survival of Human Follicular Lymphoma Cells. Cancer Discov. 2017;7(5):506–21.

27. Thomas M, Calamito M, Srivastava B, Maillard I, Pear WS, Allman D. Notch activity synergizes with B-cell-receptor and CD40 signaling to enhance B-cell activation. Blood. 2007;109(8):3342–50.

28. Ochiai K, Maienschein-Cline M, Simonetti G, Chen J, Rosenthal R, Brink R, et al. Transcriptional regulation of germinal center B and plasma cell fates by dynamical control of IRF4. Immunity. 2013;38(5):918–29.

29. Basso K, Dalla-Favera R. Roles of BCL6 in normal and transformed germinal center B cells. Immunol Rev. 2012;247(1):172–83.

30. Cinamon G, Zachariah MA, Lam OM, Foss FW, Jr., Cyster JG. Follicular shuttling of marginal zone B cells facilitates antigen transport. Nat Immunol. 2008;9(1):54–62.

31. Simonetti G, Carette A, Silva K, Wang H, De Silva NS, Heise N, et al. IRF4 controls the positioning of mature B cells in the lymphoid microenvironments by regulating NOTCH2 expression and activity. J Exp Med. 2013;210(13):2887–902.

32. Rodda LB, Bannard O, Ludewig B, Nagasawa T, Cyster JG. Phenotypic and Morphological Properties of Germinal Center Dark Zone Cxcl12-Expressing Reticular Cells. J Immunol. 2015;195(10):4781–91.

33. Santos MA, Sarmento LM, Rebelo M, Doce AA, Maillard I, Dumortier A, et al. Notch1 engagement by Delta-like-1 promotes differentiation of B lymphocytes to antibody-secreting cells. Proc Natl Acad Sci U S A. 2007;104(39):15454–9.

34. Yoon SO, Zhang X, Berner P, Blom B, Choi YS. Notch ligands expressed by follicular dendritic cells protect germinal center B cells from apoptosis. J Immunol. 2009;183(1):352–8.

35. Liu YJ, Oldfield S, MacLennan IC. Memory B cells in T cell-dependent antibody responses colonize the splenic marginal zones. Eur J Immunol. 1988;18(3):355–62.

36. Shih TA, Roederer M, Nussenzweig MC. Role of antigen receptor affinity in T cell-independent antibody responses in vivo. Nat Immunol. 2002;3(4):399–406.

37. Obukhanych TV, Nussenzweig MC. T-independent type II immune responses generate memory B cells. J Exp Med. 2006;203(2):305–10.

38. Pape KA, Kouskoff V, Nemazee D, Tang HL, Cyster JG, Tze LE, et al. Visualization of the genesis and fate of isotype-switched B cells during a primary immune response. J Exp Med. 2003;197(12):1677–87.

39. White HN, Meng QH. Recruitment of a distinct but related set of VH sequences into the murine CD21hi/CD23-marginal zone B cell repertoire to that seen in the class-switched antibody response. J Immunol. 2012;188(1):287–93.

40. Hendricks J, Visser A, Dammers PM, Burgerhof JGM, Bos NA, Kroese FGM. The formation of mutated IgM memory B cells in rat splenic marginal zones is an antigen dependent process. PLoS One. 2019;14(9):e0220933.

41. Weng AP, Millholland JM, Yashiro-Ohtani Y, Arcangeli ML, Lau A, Wai C, et al. c-Myc is an important direct target of Notch1 in T-cell acute lymphoblastic leukemia/lymphoma. Genes Dev. 2006;20(15):2096–109.

42. Chan SM, Weng AP, Tibshirani R, Aster JC, Utz PJ. Notch signals positively regulate activity of the mTOR pathway in T-cell acute lymphoblastic leukemia. Blood. 2007;110(1):278–86.

43. Gaudette BT, Roman CJ, Ochoa TA, Gomez Atria D, Jones DD, Siebel CW, et al. Resting innate-like B cells leverage sustained Notch2/mTORC1 signaling to achieve rapid and mitosis-independent plasma cell differentiation. J Clin Invest. 2021;131(20).

44. Limon JJ, Fruman DA. Akt and mTOR in B Cell Activation and Differentiation. Front Immunol. 2012;3:228.

45. Aiba Y, Kometani K, Hamadate M, Moriyama S, Sakaue-Sawano A, Tomura M, et al. Preferential localization of IgG memory B cells adjacent to contracted germinal centers. Proc Natl Acad Sci U S A. 2010;107(27):12192–7.

46. Anderson SM, Tomayko MM, Ahuja A, Haberman AM, Shlomchik MJ. New markers for murine memory B cells that define mutated and unmutated subsets. J Exp Med. 2007;204(9):2103–14.

47. Kibler A, Budeus B, Homp E, Bronischewski K, Berg V, Sellmann L, et al. Systematic memory B cell archiving and random display shape the human splenic marginal zone throughout life. J Exp Med. 2021;218(4).

48. Siu JHY, Pitcher MJ, Tull TJ, Velounias RL, Guesdon W, Montorsi L, et al. Two subsets of human marginal zone B cells resolved by global analysis of lymphoid tissues and blood. Sci Immunol. 2022;7(69):eabm9060.

49. Sperling SA, Fiedler P, Lechner M, Pollithy A, Ehrenberg S, Schiefer AI, et al. Chronic CD30-signaling in B cells results in lymphomagenesis by driving the expansion of plasmablasts and B1 cells. Blood. 2019.

50. Scheffler L, Feicht S, Babushku T, Kuhn LB, Ehrenberg S, Frankenberger S, et al. ERK phosphorylation is RAF independent in naive and activated B cells but RAF dependent in plasma cell differentiation. Sci Signal. 2021;14(682).

