## Supplementary Information for "Notch2 signaling guides B cells away from germinal centers towards marginal zone B cell and plasma cell differentiation"

### Supplementary Figures

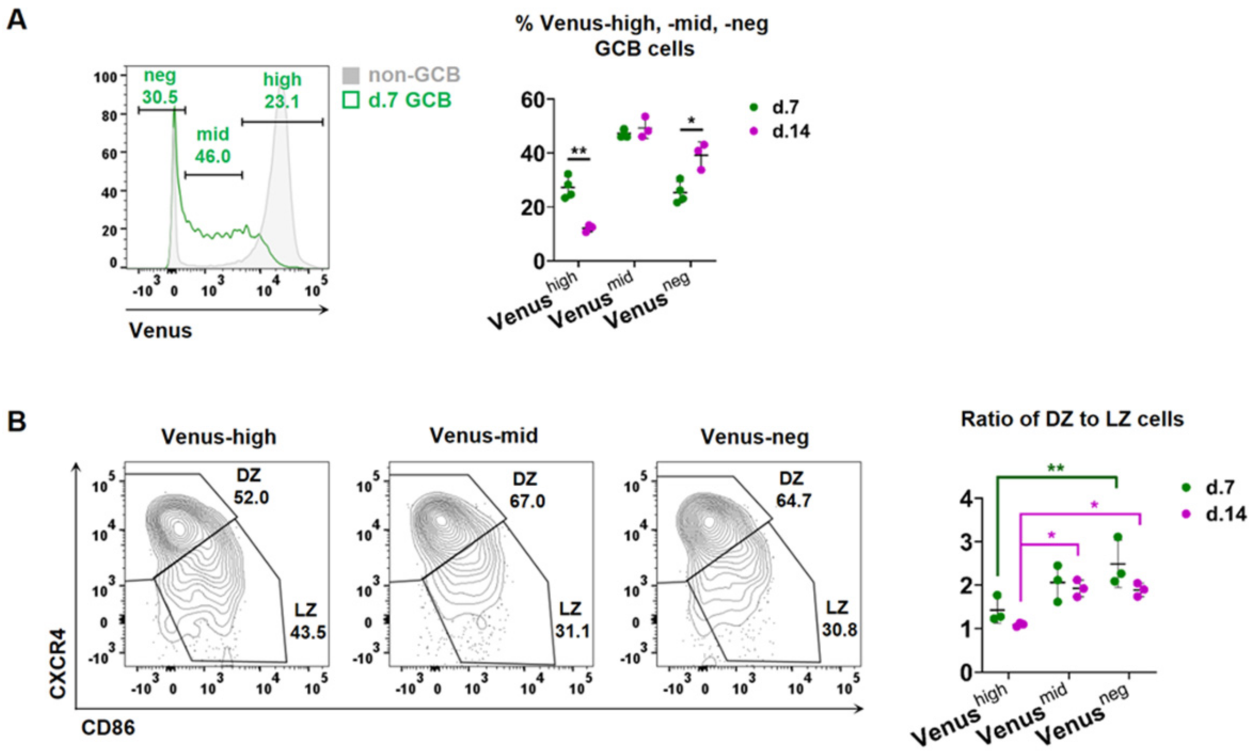


**Supplementary Fig. 1. Venus expression in the LZ and DZ of splenic GCB cells,** related to Figure 2. **(A)** The histogram shows the overlay of Venus expression in GCB and non-GCB cells at day 7 p.i. (gated as in Fig. 2A). GCB cells (CD95$^{+}$CD38$^{low}$) were divided into 3 subpopulations according to their Venus expression: Venus$^{neg}$, Venus$^{mid}$ and Venus$^{high}$. The graph depicts the percentages of the 3 subpopulations at indicated time points. **p=0.007 and *p=0.049, 2-way-ANOVA, Sidak's multiple comparisons test. d7 n=4, d14 n=3. **(B)** The Venus$^{high}$, Venus$^{mid}$, and Venus$^{low}$ GCB cells were analyzed for their DZ (CXCR4$^{high}$CD86$^{low}$) and LZ (CXCR4$^{low}$CD86$^{high}$) distribution. FACS plots are representative for day 7 p.i.. The graph depicts the ratio of the percentages of DZ to LZ cells for the 3 Venus subpopulations. A lower DZ to LZ ratio suggests an enrichment of the indicated Venus subpopulation in the LZ. **p=0.004 and *p ≤ 0.030, 2-way-ANOVA, Sidak's multiple comparisons test. n=3 mice per time point. p.i. = post immunization.

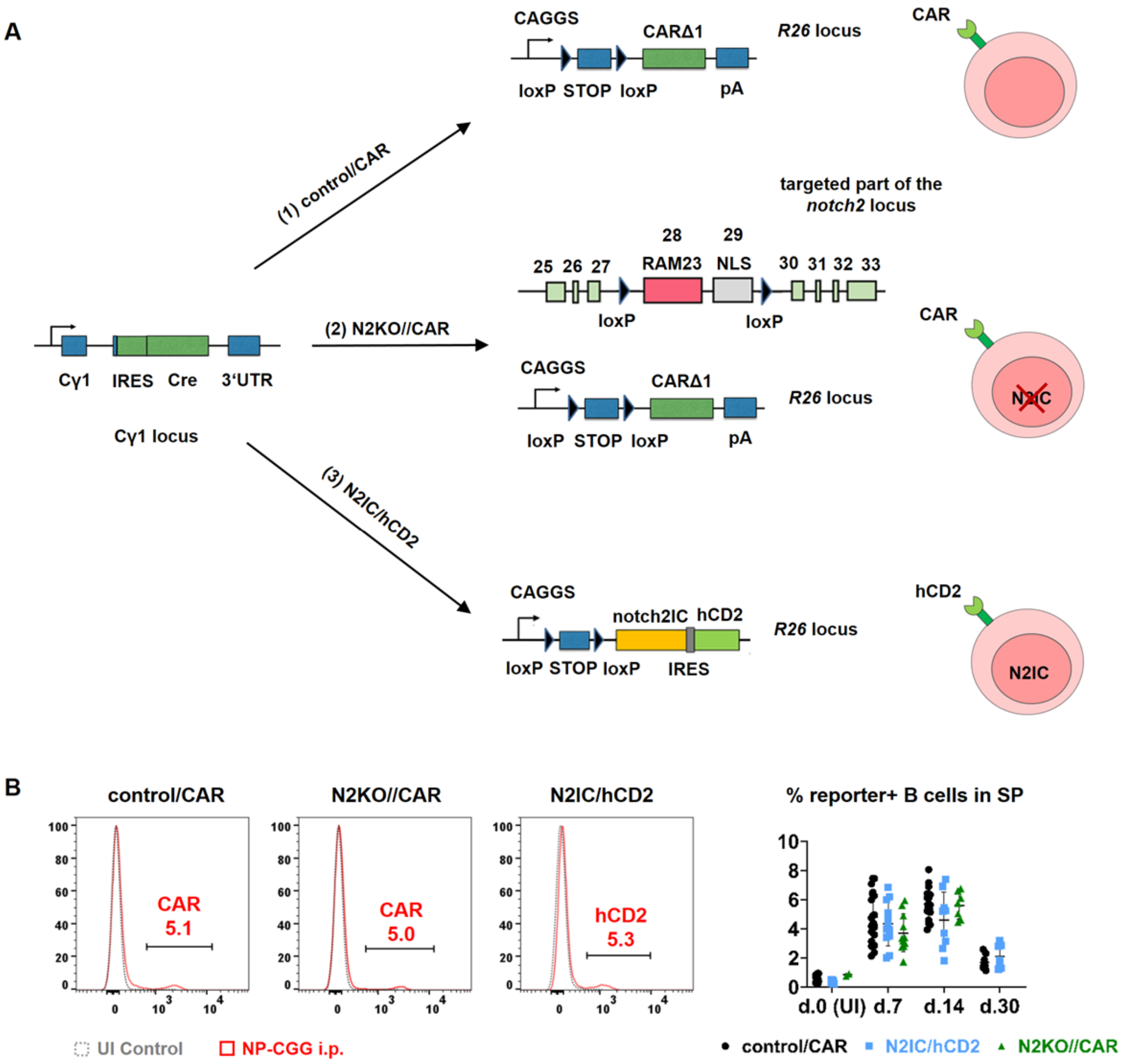


**Supplementary Fig. 2. Mouse models for antigen-dependent inactivation or constitutive activation of Notch2. (A)** To ablate or induce Notch2 upon TD immunization conditional Notch2$^{fl/fl}$ (Besseyrias et al., *JEM* 2007) or Notch2IC$^{stopfl}$ (Hampel et al., *Blood* 2011) mice were crossed with the Cγ1-Cre strain (Casola et al., *PNAS* 2006). As controls, we used the reporter mice R26-CARΔ1$^{stopF}$ (Heger et al., *Eur. J. Immunol.* 2015), which contain the coding sequence for the coxsackie/adenovirus receptor (CAR) in the *rosa26* locus. (1) Control/CAR: Cre recombinase activity leads to deletion of the stop-cassette upstream of the *carΔ1* transgene, resulting in expression of a truncated version of CAR on the cell surface. (2) N2KO//CAR: To detect cells, which deleted the Notch2-allele, we combined Notch2$^{fl/fl}$ mice with the reporter control/CAR strain. In the targeted part of the *notch2* locus, exons 28 and 29 coding for the intracellular RAM23 domain and nuclear localization signal, respectively, are flanked with loxP sites. Upon Cre-mediated recombination, both the floxed *notch2* exons and the stop-cassette upstream of *carΔ1* in the *rosa26* locus are excised, resulting in the inactivation of Notch2 (N2KO; the succeeding exons 30 to 33 of *notch2* are out of frame and are thus not translated) and expression of the CAR reporter on the cell surface. (3) N2IC/hCD2: the coding sequence for *notch2IC* is inserted into the *rosa26* locus. An IRES element and the transgene *hcd2* coding for a truncated version of the human CD2 receptor are inserted downstream of the *notch2IC*. Cre-mediated excision results in expression of Notch2IC inside the cell and hCD2 on the cell surface. **(B)** Determination of reporter gene expression by FACS analyses of splenic B lymphocytes after immunization with 100 µg NP-CGG. Histograms are pre-gated on living B220$^+$ lymphocytes**.** Percentages of reporter$^+$ B220$^+$ cells at indicated time points are summarized in the graph. Control/CAR: d0 n=11, d7 n=23, d14 n=20, d30 n=8; N2KO//CAR: d0 n=2, d7 n=11, d14 n=7; N2IC/hCD2: d0 n=5, d7 n=12, d14 n=9, d30 n=7. UI = unimmunized; NP-CGG i.p. = intraperitoneal NP-CGG injection.

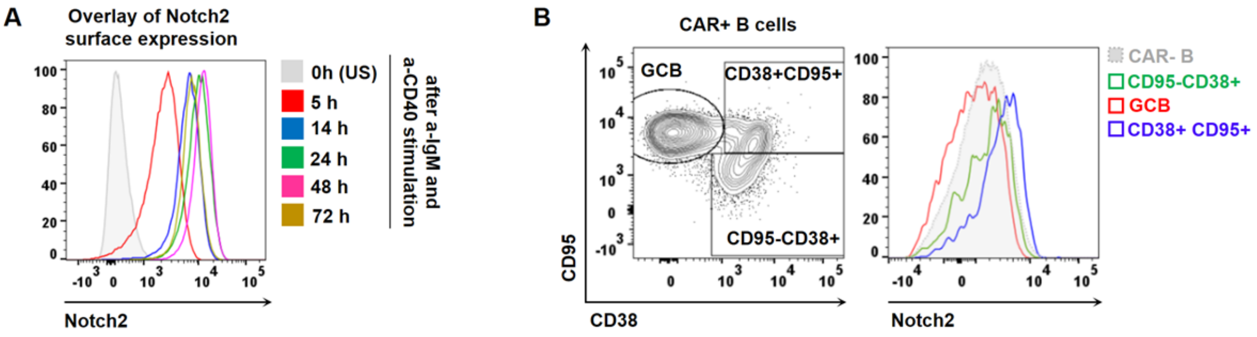


**Supplementary Fig. 3. Notch2 surface expression on FoB cells after *in vitro* stimulation and on CAR$^+$ CD38/CD95 subpopulations after TD immunization *in vivo*,** related to Figure 3. **(A)** An exemplary FACS overlay, depicting the Notch2 receptor expression on the cell surface of isolated FoB cells at indicated time points after *in vitro* stimulation with anti-CD40 and anti-IgM (0 h, unstimulated (US)). Analysis is representative for n=5 mice. **(B)** Representative FACS analysis of the gating strategy for CAR$^+$ CD38/CD95 B cell populations and a histogram overlay of the Notch2 cell surface expression on the indicated cell subpopulations at day 14 p.i.. Plots are representative for n=6 mice at day 14 p.i.. US = unstimulated cells; p.i. = post immunization.

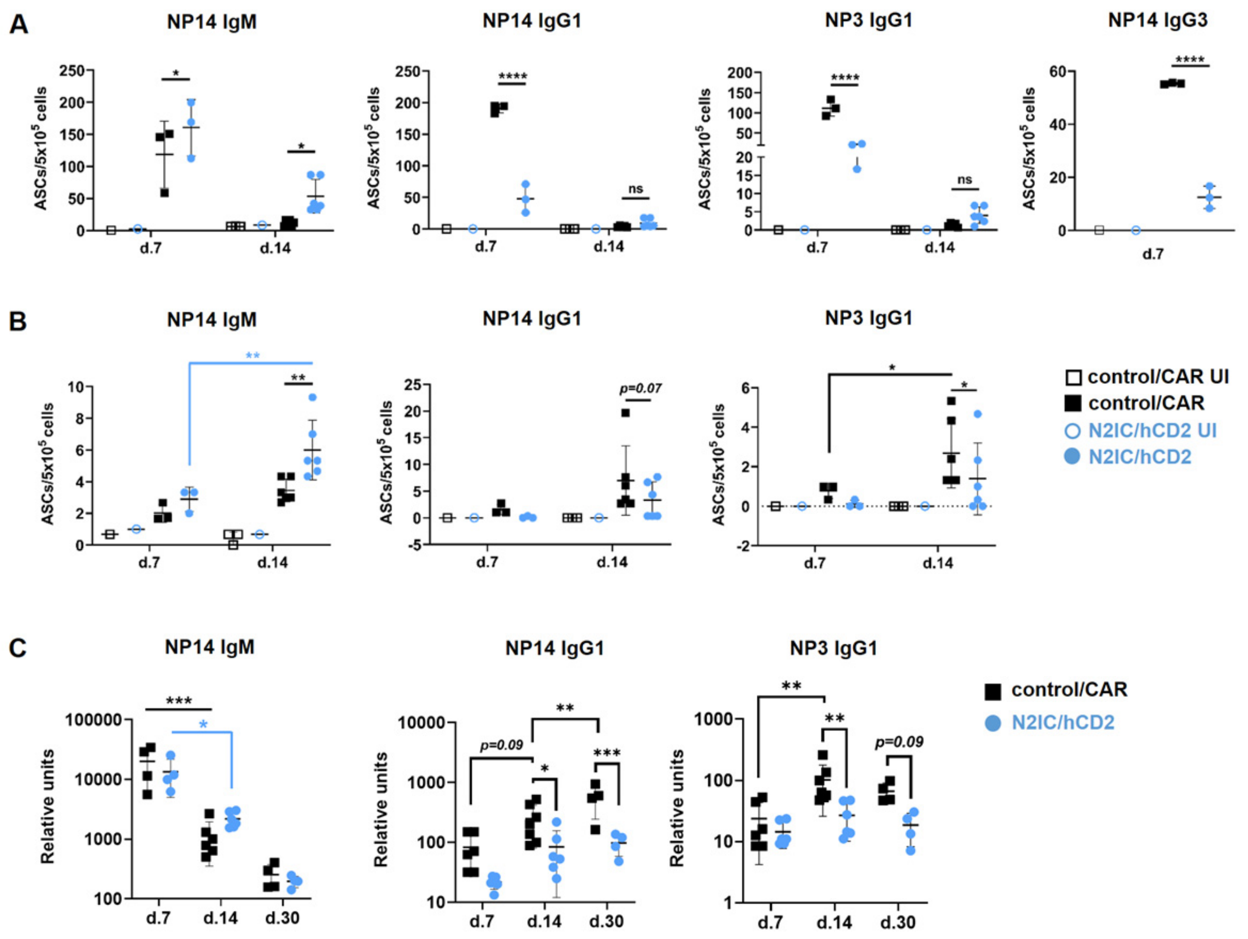


**Supplementary Fig. 4. NP-specific serum titers and NP-specific ASCs in the spleen and BM,** related to Figure 4. **(A-B)** Total (binding to NP14-BSA) and high affinity (binding to NP3-BSA) antibody-secreting cells (ASCs) of the indicated isotypes were determined in the spleen **(A)** and BM **(B)** of mice at days 7 and 14 p.i. by ELISpot analyses. Control/CAR UI: d7 n=1, d14 n=3; Control/CAR: d7 n=3, d14 n=6; N2IC/hCD2 UI: d7 n=1, d14 n=1; N2IC/hCD2: d7 n=3, d14 n=6. *p=0.05 **p=0.006 and ****p<0.0001, 2-way-ANOVA, Sidak's multiple comparisons test. **(C)** Titers of total (NP14) and high affinity (NP3) NP-specific antibodies of the indicated isotypes were measured by ELISA in the serum of N2IC/hCD2 and control/CAR mice at the indicated time points p.i.. Control/CAR: d7 n=6, d14 n=7, d30 n=4; N2IC/hCD2: d7 n=5, d14 n=6, d30 n=4. *p ≤ 0.049, **p=0.003 and ***p=0.0002, 2-way-ANOVA, Tukey's (NP14-IgM) and Sidak's (NP14-IgG1, NP3-IgG1) multiple comparisons test. UI = unimmunized; p.i = post immunization.

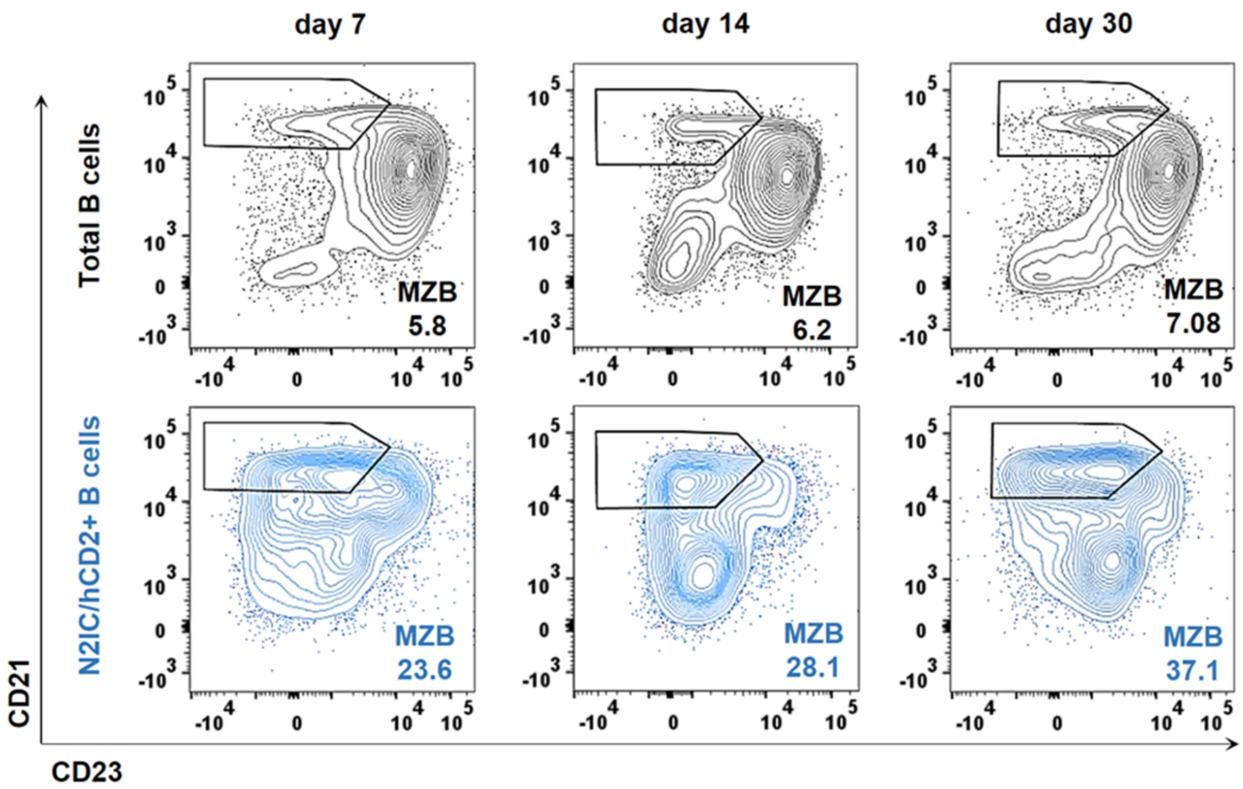


**Supplementary Fig. 5. CD23/CD21 staining to separate and quantify MZB cells in N2IC/hCD2 mice,** related to Figure 5. Representative FACS plots of the separation of $CD21^{high}CD23^{low}$ MZB cells among total $B220^{+}$ B cells (upper row, black) and $reporter^{+}$ B cells (lower row, blue) in N2IC/hCD2 mice at the indicated time points post immunization (p.i.). The analyses are representative for n=12 at day 7, n=9 at day 14 and n=7 at day 30.

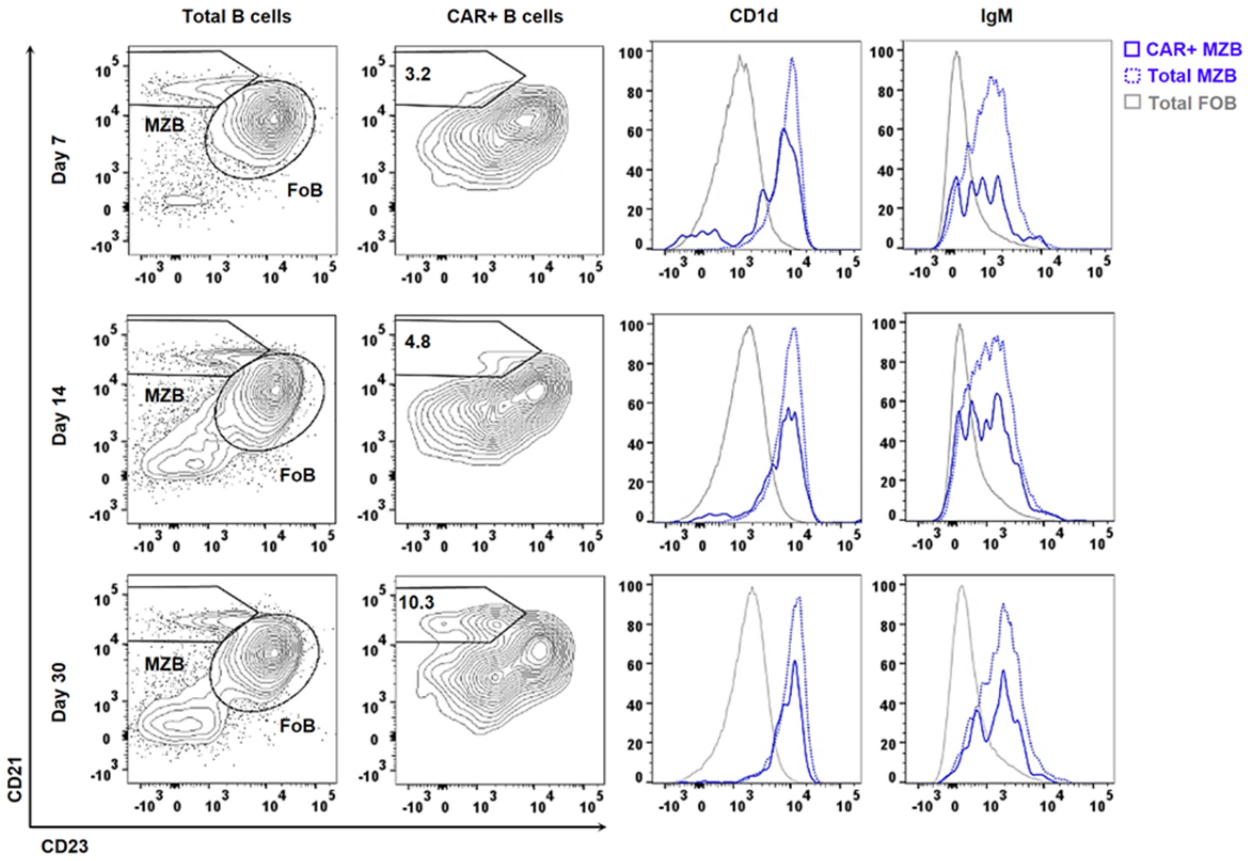


**Supplementary Fig. 6. IgM and CD1d surface phenotyping of CAR$^+$ CD23/CD21 subpopulations of control/CAR animals,** related to Figure 6. Exemplary FACS plots to illustrate the gating of total MZB cells (CD23$^{low}$CD21$^{high}$), total FoB cells (CD23$^+$CD21$^{low}$) and newly generated MZB cells (CAR$^+$CD23$^{low}$CD21$^{high}$). The histogram overlays show cell surface expression levels of CD1d and IgM on the gated CAR$^+$ MZB cell population (blue, solid), compared to the total MZB (blue, dotted) and total FoB cells (grey, solid) in control/CAR mice at indicated time points post TD-immunization with NP-CGG (p.i.). The analyses are representative for n=16 at day 7, n=17 at day 14 and n=8 at day 30.

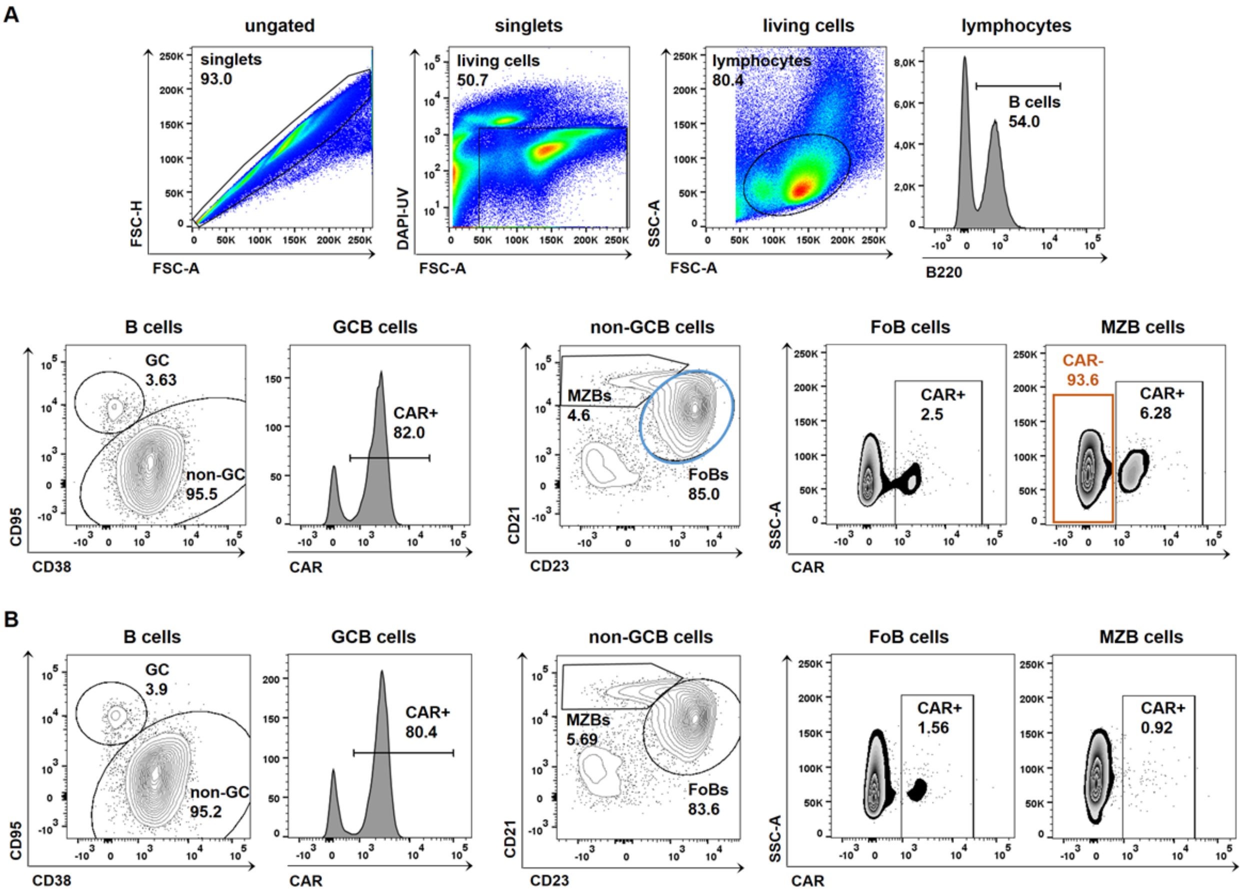


**Supplementary Fig. 7. Sequential gating for CAR-expressing cells in different B cell subpopulations for mathematical modelling,** related to Figure 7. **(A-B)** FACS plots are representative for day 7 post TD immunization with NP-CGG (p.i.). **(A)** Hierarchical gating of splenocytes in control/CAR mice to retrieve the amounts (%) of total living B220$^{+}$ B cells, GCB (CD38$^{low}$CD95$^{high}$) and non-GCB cells (CD38$^{+}$CD95$^{low}$). The frequencies of CAR$^{+}$ B cells in the GC were gated in histograms. The non-GC B cell fraction was further subdivided into MZB (CD23$^{low}$CD21$^{high}$) and FoB cells (CD23$^{+}$CD21$^{low}$). Within the non-GC MZB and FoB cells, the percentages of CAR-expressing cells were gated in CAR vs. SSC-A plots. The frequencies of CAR$^{-}$ cells within the non-GC MZB cell fraction (orange) and the total non-GC FoB cells (blue) were used for generating the plots in Supplementary Fig. 8 below. The gating strategy is representative for n=3 at day 4, n=9 at day 7, n=3 at day 9, n=15 at day 14, n=3 at day 17, n=5 at day 22, n=4 at day 26, n=6 at day 30 post immunization. **(B)** Exemplary analysis of splenocytes from N2KO//CAR mice. Singlets, living cells, lymphocytes, reporter+ GCB cells and reporter+ non-GC FoB and MZB cells were sequentially gated as in (A). The gating strategy is representative for n=7 at day 7, n=5 at day 9, n=6 at day 14 post immunization.

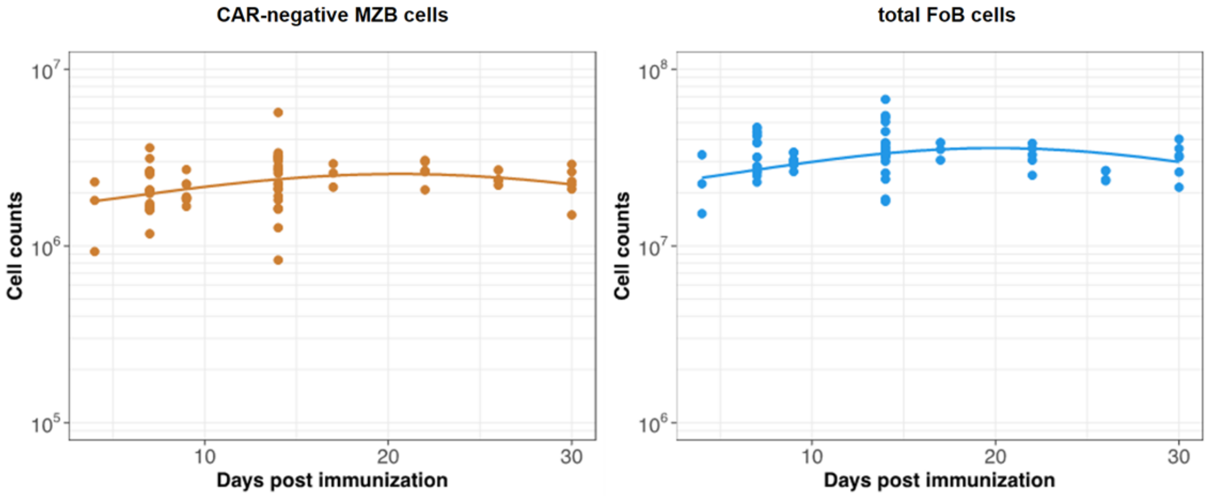


**Supplementary Fig. 8. Empirical descriptions of the dynamics of the numbers of FoB and MZB cells,** related to Figure 8. We show here curves for total FoB cells and $CAR^-$ MZB cells using the functions (defined in Supplementary Text 1), which were used as inputs in our ODE model systems. The y-axis depicts the total cell numbers of the respective cell population. d4 n=3, d7 n=9, d9 n=3, d14 n=15, d17 n=3, d22 n=5, d26 n=4, d30 n=6 control/CAR mice following TD-immunization.

### Supplementary Tables

#### Supplementary Table 1. List of antibodies used in this study.

| | Antibody | Conjugate | Clone | Company | Dilution factor |
|---|---|---|---|---|---|
| FACS | CD1d | Alexa Fluor 647 | 1B1 | BioLegend | 1:200 |
| | TACI | APC | 8G10-3 | eBioscience | 1:100 |
| | CD86 | APC | / | BD Biosciences | 1:100 |
| | CAR | FITC | sc-56892 | Santa Cruz | 1:25 |
| | hCD2 | FITC | REA972 | miltenyi | 1:50 |
| | hCD2 | PeVio770 | / | miltenyi | 1:100 |
| | hCD2 | APC; PE | RPA-2.10 | eBioscience | 1:100; 1:100 |
| | CXCR4 (CD184) | PE | 2B11 | eBioscience | 1:400 |
| | CD38 | PeVio770 | / | miltenyi | 1:100 |
| | B220 | PerCP; APC; FITC; PE | RA3-6B2 | BD Biosciences | 1:100; 1:250; 1:200; 1:350 |
| | CD21 | BV421 | 7G6 | BD Biosciences | 1:200 |
| | CD43 | Biotin; BV421 | S7 | BD Biosciences | 1:350; 1:200 |
| | Streptavidin (SA) | PerCP; APC | | BD Biosciences | 1:100; 1:400 |
| | CD23 | PE; Alexa Fluor 647; PeVio770 | B3B4 | BD Biosciences | 1:200; 1:100; 1:200 |
| | CD5 | BV421 | 53-7.3 | BD Biosciences | 1:200 |
| | IgM | APC | II/41 | BD Biosciences | 1:100 |
| | IgM | BV421; PeVio770 | R6-60.2 | BD Biosciences | 1:100; 1:300 |
| | AA4.1 | PE | AA4.1 | eBioscience | 1:200 |
| | CD95 | BV421 | JO2 | BD Biosciences | 1:200 |
| | CD138 | BV421 | 281-2 | BD Biosciences | 1:200 |
| | Blimp1 | PE | sc47732 PE | Santa Cruz | 1:50 |
| | Bcl6 | Alexa Fluor 647 | K112-91 | BD Biosciences | 1:50 |
| | Irf4 | eFluor660; PE | 3E4 | eBioscience | 1:100; 1:500 |
| | CD19 | PeVio770 | REA749 | miltenyi | 1:200 |
| | CD19 | BV510 | 1D3 | BD Biosciences | 1:200 |

| | | | | | |
|---|---|---|---|---|---|
| | Notch2 | PE | HMN2-35 | BioLegend | 1:180 |
| | IgD | APC | 11-26c.2a | BD Biosciences | 1:300 |
| ELISpot | rat-anti-mouse IgM | Biotin | R6-60.2 | BD Biosciences | 1:500 |
| | rat-anti-mouse IgG1 | Biotin | A85-1 | BD Biosciences | 1:500 |
| | rat-anti-mouse IgG3 | Biotin | R40-82 | BD Biosciences | 1:500 |
| | Avidin D | HRP | | Vector | 1:2000 |
| ELISA | rat-anti-mouse IgM | HRP | YF97 | Southern Biotech | 1:5000 |
| | rat-anti-mouse IgG1 | Biotin | A85-1 | BD Biosciences | 1:500 |
| | Avidin D | HRP | | Vector | 1:2000 |
| Histology | Rat-anti-mouse-hCD2 | Biotin | RPA-2.10 | BD Biosciences | 1:50 |
| | Rabbit-anti-mouse Laminin | | L9393 | Sigma-Aldrich | 1:100 |
| | Streptavidin | Alkaline Phosphatase | | Sigma-Aldrich | 1:200 |
| | anti-Rabbit IgG | Peroxidase | | Sigma-Aldrich | 1:200 |
| | Rat-anti-mouse CD90.2 (Thy1.2) | Biotin | 53-2.1 | BD Biosciences | 1:100 |
| | Rat-anti-mouse B220 | APC | RA3-6B2 | BD Biosciences | 1:500 |
| | Rat-anti-mouse Irf4 | | 3E4 | eBioscience | 1:100 |
| | Goat-anti-rabbit IgG | Cyanine Cy3 | | Jackson ImmunoResearch | 1:500 |
| | Goat-anti-rat IgG | Alexa Fluor 488 | | Jackson ImmunoResearch | 1:500 |
| | Streptavidin | Alexa Fluor 594 | | Life Technologies | 1:500 |
| | Rat-anti-mouse GL-7 | FITC | GL7 | BD Biosciences | 1:100 |
| | Rat-anti-mouse GL7 | Alexa Fluor 647 | GL7 | BioLegend | 1:100 |
| | Rat-anti-mouse CD19 | APC | 1D3 | BD Biosciences | 1:200 |
| | Rat-anti-mouse MOMA1 | Biotin | ab51814 | abcam | 1:100 |

### Supplementary Notes

**Supplementary Text 1. Defining phenomenological functions to capture the dynamics of precursor populations for the GCB and CAR-expressing MZB cells**

**Dynamics of FoB cells:** In the branched model, we assumed that total FoB cells differentiate into MZB cells. We used the following empirical descriptor function to capture the time course of counts of total FoB cells.

$$\phi(t) = \phi_0 \ (1 + e^{-\nu\,(t-b_0)^2}) \qquad \text{Supplementary equation (1)}$$

We estimated the parameters $\phi_0, \nu$ and $b_0$ by fitting the supplementary equation (1) to the log-transformed numbers of FoB cells. The fit is shown in the Supplementary Fig. 8.

**Dynamics of CAR$^-$ MZB cells:** In all our models we considered that activation of CAR$^-$ MZB cells results in CAR upregulation. The time course of total numbers of CAR$^-$ MZB cells was also modeled using the supplementary equation (1) and the corresponding model fit is shown in the Supplementary Fig. 8.

**Supplementary Text 2. Model ranking and selection criteria**

We fitted each model ($M_j$) described in **Materials and Methods**, section ***Mathematical models of B cell dynamics during a TD immune response*** simultaneously to the time-courses of cell counts of CAR$^+$ MZB cells, CAR$^+$ GCB cells and total B cells from control/CAR and N2KO//CAR mice. The model parameters (expressed as a vector $\theta$) and the errors associated with the empirical measurements in each dataset ($\sigma_i$) are then estimated using the Bayesian statistical inference approach. We assumed that the residuals are normally distributed. The likelihood and the probability density for each dataset are then defined as,

$$\begin{aligned} y_i &\sim \text{Normal}(\mu_i, \sigma_i) && \text{[Likelihood]} \\ \mu_i &= M_j(\text{time}_i, \theta) && \text{[Model prediction]} \\ P(y_i|\theta) &= \frac{1}{2\pi\,\sigma^2}\exp\left(-\frac{(y_i-\mu_i)^2}{2\,\sigma^2}\right) && \text{[Probability density]} \end{aligned}$$

Supplementary Equation (2)

We defined the 'prior' distributions of model parameters representing our assumptions regarding their values. The Bayesian procedure then updates the priors using the likelihood, to generate posterior distribution ($\hat{\theta}$) of model parameters conditional on the evidence from the data. The input for this procedure is the joint probability densities of three datasets.

$$P(\hat{\theta}|y_1, y_2, y_3) = \frac{P(y_1|\hat{\theta}) \cdot P(y_2|\hat{\theta}) \cdot P(y_3|\hat{\theta}) \cdot P(\theta)}{P(y_1) \cdot P(y_2) \cdot P(y_3)}$$

Supplementary Equation (3)

Models are fitted using the no-U-turn-sampler (*NUTS*) sampler in the Stan programming language, where Parameters are sampled from the joint density $P(\theta)$ following Hamiltonian Monte Carlo algorithm (Stan Development Team, 2022). The descriptions of model systems, the prior distributions of parameters and the likelihood definitions are encoded in the Stan language and are available at the linked Github repository.

We compared the support for models using the leave-one-out (LOO) cross validation method, which estimates the expected log point-wise predictive density (*elpd*) for each model $M_j$ – the measure of its out-of-sample prediction accuracy (Vehatari et al., Statistics and Computing 2016). We estimate the probability density of $y_i$ given the model $M_j$ fitted on the data with observation $i$ excluded → $P(y_i|y_{-i},\ M_j)$. This leave-one-out process is repeated for all $'n'$ observations in the data. The *elpd* estimate and its standard error are then calculated as,

$$\widehat{\text{elpd}}^{j}_{\text{loo}} = \sum_{i=1}^{n} \text{elpd}^{j}_{\text{loo, i}} = \sum_{i=1}^{n} \log(P(y_i|y_{-i}, M_j),$$

$$\text{se}(\widehat{\text{elpd}^{k}_{\text{loo}}}) = \sqrt{\sum_{i=1}^{n} \left(\text{elpd}^{k}_{\text{loo, i}} - \text{elpd}^{k}_{\text{loo}}/n\right)^2}.$$

Supplementary Equation (4)

We used the *loo-2.0* package in the *Rstan* library to estimated *elpd*, which employs Pareto smoothed importance sampling (PSIS) (Vehatari et al., Statistics and Computing 2015) to approximate LOO cross validation. The estimates of elpd and its standard error were used to rank models using the Pseudo-Bayesian model averaging (BMA) method (Yao et al., 2018) implemented in the *loo-2.0* package. The model weight implies relative support for each model, analogous to the model weights calculated using Akaike's Information Criterion (AIC) (Akaike, 1978; Burnham, 2002; Wagenmakers and Farrell, 2004), and is given as,

$$W_k = \frac{\exp\left(\widehat{\text{elpd}^{k}_{\text{loo}}} - \frac{1}{2}\text{se}(\widehat{\text{elpd}^{k}_{\text{loo}}})\right)}{\sum_{k=1}^{K} \exp\left(\widehat{\text{elpd}^{k}_{\text{loo}}} - \frac{1}{2}\text{se}(\widehat{\text{elpd}^{k}_{\text{loo}}})\right)}.$$

Supplementary Equation (5)

As the models' weight calculations are based on the *elpd* estimates from LOO cross-validation, it also denotes the confidence in model's ability to predict new data relative to all the other models considered in the analysis. Model weights for all the fitted models are shown in **Table 1** of the Results in the main text.

### Supplementary References


Besseyrias V, Fiorini E, Strobl LJ, Zimber-Strobl U, Dumortier A, Koch U, et al. Hierarchy of Notch-Delta interactions promoting T cell lineage commitment and maturation. J Exp Med. 2007;204(2):331-43.

Hampel F, Ehrenberg S, Hojer C, Draeseke A, Marschall-Schroter G, Kuhn R, et al. CD19-independent instruction of murine marginal zone B-cell development by constitutive Notch2 signaling. Blood. 2011;118(24):6321-31.

Casola S, Cattoretti G, Uyttersprot N, Koralov SB, Seagal J, Hao Z, et al. Tracking germinal center B cells expressing germ-line immunoglobulin gamma1 transcripts by conditional gene targeting. Proc Natl Acad Sci U S A. 2006;103(19):7396-401.

Heger K, Kober M, Riess D, Drees C, de Vries I, Bertossi A, et al. A novel Cre recombinase reporter mouse strain facilitates selective and efficient infection of primary immune cells with adenoviral vectors. Eur J Immunol. 2015;45(6):1614-20.

Stan Development Team (2022) Stan modeling language user's guide and reference manual, version 2.29 Stan.
https://mc-stan.org

Vehtari A, Gelman A, Gabry J. Practical Bayesian model evaluation using leave-one-out cross-validation and WAIC, Journal: Statistics and Computing, 2016 doi: 10.1007/s11222-016-9696-4

Vehtari A, Gelman A, Gabry J. Efficient implementation of leave-one-out cross-validation and WAIC for evaluating fitted Bayesian models, Journal: Statistics and Computing 2015 arXiv:1507.04544v1

Yao Y, Vehtari A, Simpson D, Gelman A. Using Stacking to Average Bayesian Predictive Distributions, Journal Bayesian Analysis 2018 https://doi.org/10.1214%2F17-ba1091

Hirotugu Akaike. On the Likelihood of a Time Series Model, Journal of the Royal Statistical Society 1978. Series D (The Statistician) Vol. 27, No. 3/4, https://doi.org/10.2307/2988185

Burnham KP, Anderson DR. , Model Selection and Multimodel Inference: A Practical Information-Theoretic Approach, 2002 Springer-Verlag, arXiv:1011.1669v3

Wagenmakers EJ, Farrell S. AIC model selection using Akaike weights, Psychon Bull Rev, 2004 DOI: 10.3758/bf03206482
